# Microglial colonisation of the developing brain is facilitated by clonal expansion of highly proliferative progenitors and follows an allometric scaling

**DOI:** 10.1101/2022.09.15.507569

**Authors:** Liam Barry-Carroll, Philip Greulich, Abigail R. Marshall, Kristoffer Riecken, Boris Fehse, Katharine E. Askew, Kaizhen Li, Olga Garaschuk, David A. Menassa, Diego Gomez-Nicola

## Abstract

Microglia are the resident immune cells of the brain and arise from yolk sac-derived macrophages during early embryogenesis. On entering the brain, microglia undergo *in situ* proliferation and eventually colonise the entire brain by the second and third postnatal weeks in mice. However, the intricate dynamics of their developmental expansion remain unclear. Here, we examine and characterise the proliferative dynamics of microglia during embryonic and postnatal development. Using complementary fate-mapping techniques, we demonstrate that the developmental colonisation of the brain by microglia is facilitated by clonal expansion of highly proliferative microglial progenitors that occupy spatial niches throughout the brain. We also find that the distribution of microglia switches from a clustered to a random pattern between embryonic and late postnatal development. Moreover, the developmental increase in microglia follows the proportional growth of the brain in an allometric manner with the density of microglia eventually stabilising when the mosaic distribution has been established. Overall, our findings offer insight into how the competition for space acts as a driving force for microglial colonisation by clonal expansion during development.

## INTRODUCTION

Microglia, the brain’s parenchymal macrophages, primarily derive from yolk sac macrophages which are generated at embryonic day 7.5 (E7.5) in mice and appear first within the brain at around E9.5 (Alliot et al., 1999; Ginhoux et al., 2010). The number of infiltrating yolk sac macrophages is dramatically reduced following the establishment of the blood brain barrier at E14.5 (Ranawat and Masai, 2021; Stremmel et al., 2018). The entry of ‘foreign’ microglial precursors is crucial for normal brain development processes: microglial cells participate in the wiring of neuronal networks through pruning of dopaminergic axons and the establishment of the correct positioning of interneurons in their respective laminae (Squarzoni et al., 2014); they support myelination (Wlodarczyk et al., 2017) and clear myelin debris and apoptotic cells (Cunningham et al., 2013); and they promote the maturation and differentiation of neurons and oligodendrocytes (Decoeur et al., 2022; Marsters et al., 2020).

The majority of microglial cells are born during the first two postnatal weeks and this is reflected by a peak in their proliferation followed by a notable rise in cell numbers (Alliot et al., 1999; Askew and Gomez-Nicola, 2017; Askew et al., 2017; De et al., 2018; Nikodemova et al., 2015). Following the developmental expansion phase, stable densities of microglia are eventually achieved in a region-specific manner through a fine-tuned balance of proliferation and apoptosis-driven refinement of microglial numbers (Askew et al., 2017; Hope et al., 2020). There are two possibilities for how the population expands: the first comes from sequencing studies which show several clusters of microglia with a transcriptomic profile enriched in cell-cycle and proliferation-related genes suggesting a unique proliferative capacity for these cells during development (Hammond et al., 2019; Li et al., 2019; Masuda et al., 2019). Another possibility is that all microglial precursors share the same propensity to proliferate, akin to the reported kinetics of microglia in the adult brain whereby microglia proliferate in a stochastic manner that is not restricted to a progenitor niche (Askew et al., 2017; Tay et al., 2017). In contrast to this steady-state model of turnover, the microglial population is suggested to undergo context-dependent clonal expansion in response to both acute (Tay et al., 2017) and chronic insults (Jordão et al., 2019) resulting in a significant increase in the density of microglia.

Some proposed drivers of microglial proliferation during development include canonical mitogenic signalling via the CSF1/CSF1R axis (Squarzoni et al., 2014). Another potential driver of microglial proliferation is the availability of space whereby in the presence of unoccupied spatial niches, microglia will proliferate until a level of contact inhibition is achieved similar to what has been observed in depletion/repopulation paradigms (Zhan et al., 2019), although this mechanism of expansion has not been studied during development. Indeed, a hallmark of microglia in the adult brain is their ‘mosaic-like’ distribution throughout the parenchyma which allows them to survey their local environment in a territorial fashion without input from neighbouring cells (Nimmerjahn et al., 2005). However, how this mosaic distribution is achieved is yet to be uncovered and studied within the framework of microglial developmental dynamics.

In contrast to this regular adult mosaic distribution, the topographical distribution of microglia during development is heterogenous, with accumulation of microglial cells in various neuronal progenitor niches throughout the brain including the deep layers of the cortex and various axonal tracts such as the corpus callosum (Squarzoni et al., 2014). These hotspots of microglia were traditionally referred to as ‘fountains of microglia’ and were thought to represent regions of substantial infiltration of microglial precursors from the periphery (Río-Hortega, 1919). It is now appreciated that the topographical placement of microglia during development is influenced by their local functions and interactions with the cellular milieu of the developing brain. A notable example is within the embryonic forebrain where microglia transiently associate with axonal tracts of dopaminergic neurons and play a vital role in modulating proper wiring of neuronal networks at this early stage (Squarzoni et al., 2014). The extent to which these interactions dictate the final placement/colonisation of the microglial population during development remains to be seen.

While there have been a number of exciting findings in regard to the developmental kinetics of the microglial population, it still remains unclear how their expansion is orchestrated in a relatively short period and how they achieve their final mosaic-like distribution. A pivotal question is whether a highly proliferative niche of microglia exists or if each progenitor shares an equal propensity to proliferate. Here, we used a combined fate-mapping approach and applied a battery of spatial analyses to investigate whether microglia clonally expand during development and how the spatial distribution of microglia changes during development. Furthermore, we wanted to see how the concept of space-availability impacts onto the expansion of microglia.

Our findings support the notion that the early expansion of the microglial population is coupled to the growth of the brain and space availability until a stable mosaic distribution is achieved. The developmental expansion of microglia is facilitated by clonal growth of embryonic progenitors which colonise the entire parenchyma. Finally, using an RGB marking strategy, we uncover the clonal nature of the dynamics of microglia, revealing specific cells with high proliferative potential. Our data show that the relatively ‘homogenous’ mosaic of microglia in the adult brain is comprised of a diverse array of interlocking clones which differ by size and regional association.

## RESULTS

### The developmental expansion of microglia follows an allometric growth coupled to the relative change in brain area

We set out to characterise the spatiotemporal changes in microglia that are associated with development of the population. We used the lineage defining transcription factor PU.1 to map the changes in density and distribution of microglia across the embryonic forebrain, midbrain and hindbrain. All PU.1^+^ microglia (Figure 1A) were counted in these areas and overall, we observed a remarkable stability in the density of microglia between E10.5 and E18.5 in the forebrain and midbrain (Figure 1B). We also found that the magnitude of the change in the average number of microglial cells closely followed the trend of change in proportional brain area (Figure 1B), suggesting that microglial expansion is likely coupled to the growth rate of the brain. This trend was not observed in the hindbrain with the density of microglia significantly peaking during early embryonic (E10.5 and E12.5) and late embryonic development (E18.5) (Figure 1B). In contrast with embryonic development, the density of PU.1^+^ microglia rapidly increased in the postnatal forebrain and peaked at P14 (Figure 1C). By P21, there was a non-significant drop in the density of microglia in line with a reported refinement of the population (Askew et al., 2017; Hope et al., 2020; Nikodemova et al., 2015). Interestingly, this trend was not observed when analysing the average change in numbers suggesting that the increase in forebrain area may contribute to an overall decrease in density by P21, similar to what has been reported in the basal ganglia (Hope et al., 2020). Moreover, we observed that the magnitude of change in the average number of microglial cells closely followed that of the proportional change in brain area. Moreover, analysis of the proliferative index of embryonic microglia (IBA1^+^ Ki67^+^/ IBA1^+^) indicated that microglial proliferation was largely stable between E10.5 and E16.5 (Figure 1D, E). Taken together, these findings suggest that microglial expansion follows an allometric growth curve that is coupled to the changes in brain growth.

**Figure 1.**
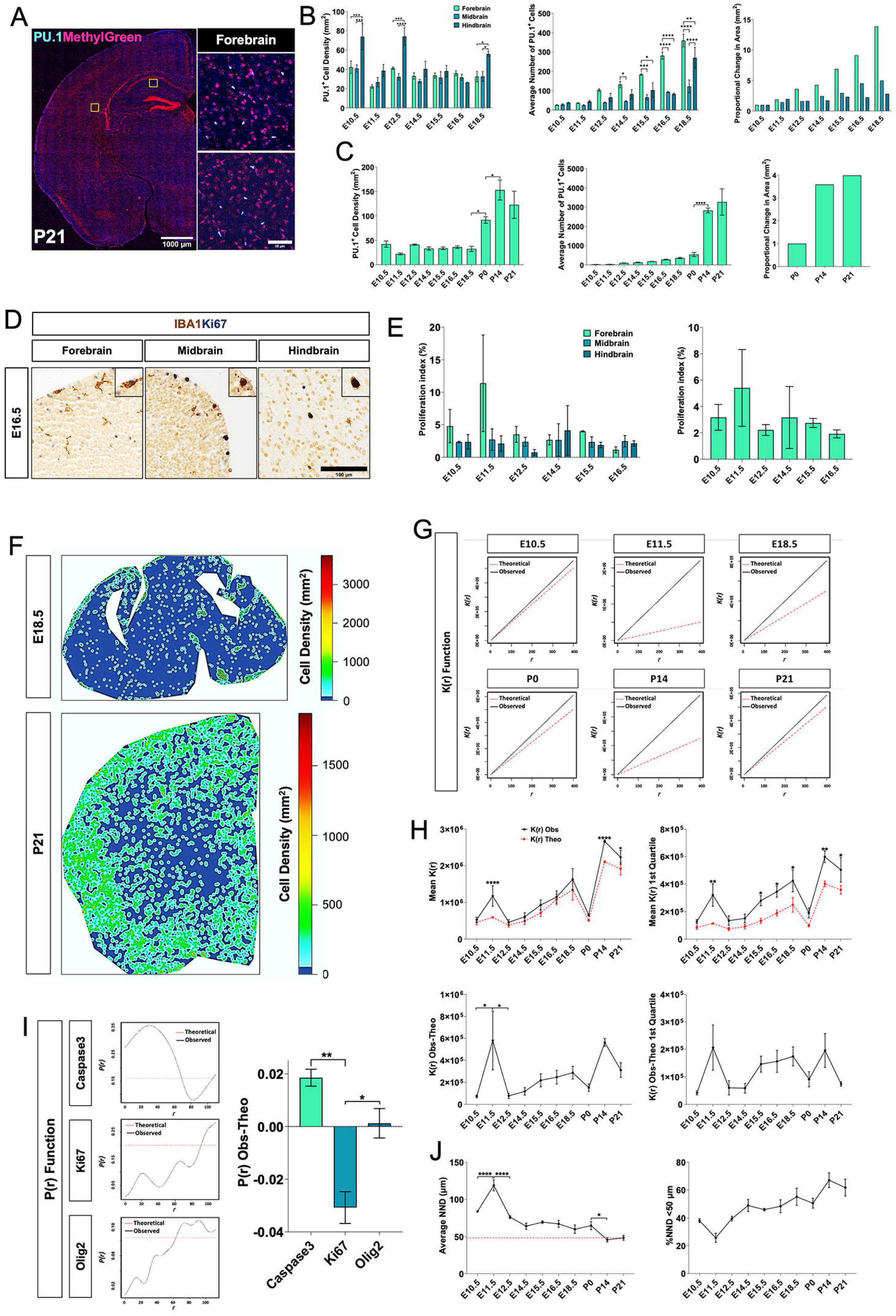
The spatiotemporal dynamics of microglia throughout development. (A) Representative image of PU.1^+^ cells at E18.5 in the forebrain, midbrain and hindbrain (images inverted for clarity). (B) Time-course analysis of the number of PU.1^+^ microglia in the forebrain, midbrain and hindbrain during embryonic development. (C) Representative image PU.1^+^ cells at P21 in the forebrain (images inverted for clarity). (D) Time-course analysis of the number of PU.1^+^ microglia in the forebrain and proportional change in area during embryonic and postnatal development. (E) Representative density maps of PU.1^+^ cells at E18.5 and P21. (F) Time-course analysis of PU.1^+^ microglia NND during embryonic and postnatal development. (G) Representative K(r) functions of PU.1^+^ microglial cells during development. (H) Ripley’s K analysis of the of PU.1^+^ microglia over the entire K(r) function and over the first quartile. (I) Normalised Ripley’s K analysis of the of PU.1^+^ microglia over the entire K(r) function and over the first quartile. Scale bars are 20 μm and 1000 μm in (A) and (B). Data are expressed as mean ± SEM, n=3 per group (1 litter per time point), Two-way ANOVA followed by Tukey’s post-hoc analysis (B and H) and One-way ANOVA followed by Bonferroni’s post-hoc analysis for pairwise comparison (D, F and I). Statistical differences: *p<0.05, **p<0.01, ***p<0.001, ****p<0.0001.

### The mosaic distribution is achieved by late postnatal development coinciding with the levelling of microglial density

Having observed that changes in microglial numbers closely follow the changes in brain size, we sought to investigate in detail the relationship between microglia density and spatial distribution. Heterogenous spatial trends were observed among embryonic microglia that were characterised by either a clustered distribution or a dispersed distribution throughout the brain (Figure 1F). In contrast to the trends observed during embryonic and early postnatal development, microglia assumed an even spatial distribution by P21 (Figure 1F). This distribution was particularly notable in the cerebral cortex and is in line with the reported mosaic or tiled distribution of microglia in the adult brain. In-depth spatial analysis was carried out using Ripley’s K function (K(r)). Briefly, the K(r) was calculated for our spatial points based on the XY coordinates of PU.1^+^ labelled microglia (observed) and compared to a set of randomly generated spatial points (theoretical) for the same window assuming a Poisson distribution. An observed K(r) value that is greater than the corresponding theoretical K(r) value indicates that those points are clustered. Conversely, a dispersed distribution is assumed when the observed K(r) is less than the corresponding theoretical K(r). Time course analysis of the microglial K(r) revealed that the spatial dynamics of microglia varies during development, with several observed shifts between a clustered to a random distribution at E11.5-E12.5, E18.5-P0 and P14-P21 (Figure 1G, H). Several factors likely contribute to these changes in spatial dynamics including the overall migratory behaviour of microglia, their specific developmental functions and inter-cellular interactions as microglia are highly receptive cells engaging in multiple forms of cell-to-cell communication.

To test whether the heterogenous spatial profile of developmental microglia is influenced by their functions and/or interactions with other cell types, the MarkConnect function was computed to investigate the spatial relationship between microglia (PU.1^+^ or IBA1^+^) and either dying cells (Caspase3^+^), proliferating (Ki67^+^) cells or OPCs (Olig2^+^) as recent studies have suggested that microglia interact with the aforementioned elements during development (Cunningham et al., 2013; Hattori et al., 2020; Marsters et al., 2020). Briefly, the MarkConnect function was used to test spatial dependence or independence between both cell types at E16.5 within the forebrain. Overall, we observed that microglia displayed a dependent relationship with Caspase3^+^ cells whereas an this relationship was not observed between microglia and Ki67^+^ cells (Figure 1I, Supplementary Figure 1). Surprisingly, we found no distinct trend between microglia and Olig2^+^ cells (Figure 1I, Supplementary Figure 1) which is in contrast to a recent study which found that microglia were localised to regions with a high density of OPCs (Marsters et al., 2020). Therefore, the cellular interactions of microglia with dying and proliferating cells may drive their dispersion and migration throughout the brain accounting for the level of cell dispersion observed in postnatal stages.

The heterogenous distribution of microglia during development is in stark contrast to their reportedly regular distribution in the adult brain which has been likened to that of a mosaic pattern characterised by an average NND of 50 µm between cells (Nimmerjahn et al., 2005). Therefore, we investigated how the NND of microglia changes during development. We found that the average NND between microglia was highest during early embryonic development at E11.5 (Figure 1J). Following E11.5, there was a steady decline in the NND as development progressed and the density of microglia increased (Figure 1J). Notably, the average NND of microglia was lowest at P14 when the percentage of microglia in close proximity to neighbouring cells (NND <50 µm) was highest at 67% coinciding with the observed peak in microglial density at P14 (Figure 1J). Moreover, changes in the NND of microglia significantly correlated with changes in microglial cell density (Supplementary Figure 2 A,B). Altogether these findings suggest that the spatial profile of microglia during early development is highly dynamic and heterogenous in contrast to P21 when the cells display an even mosaic distribution that is achieved following postnatal refinement of the population.

### Inhibition of postnatal microglial apoptosis alters the spatial distribution of microglia

We aimed to test the interdependence of microglia density and the formation of the mosaic, by using *Vav-Bcl2* mice, which we previously showed to have increased microglial density due to the overexpression of antiapoptotic protein BCL2 (Askew et al., 2017). In adult *Vav-Bcl2* mice, the density of microglia is significantly higher and the NND of microglia is significantly lower than that of microglia from wild-type mice indicating that the spatial dynamics of microglia become altered in the face of overcrowding (Figure 2 A,B). On average, microglia covered a significantly smaller territory thus accommodating an elevated cell density while still maintaining a minimal overlap between cells (Figure 2 A,B). We performed spatial analysis using the Ripley’s K function (K(r)) and observed that the changes in NND and territory do not happen at the expense of altering the distribution, as Vav-Bcl2 mice maintained a random distribution similar to wild-type mice (Figure 2C). These findings suggest that the typical mosaic distribution and spatiotemporal profile of microglia is directly influenced by space availability within the brain.

**Figure 2.**
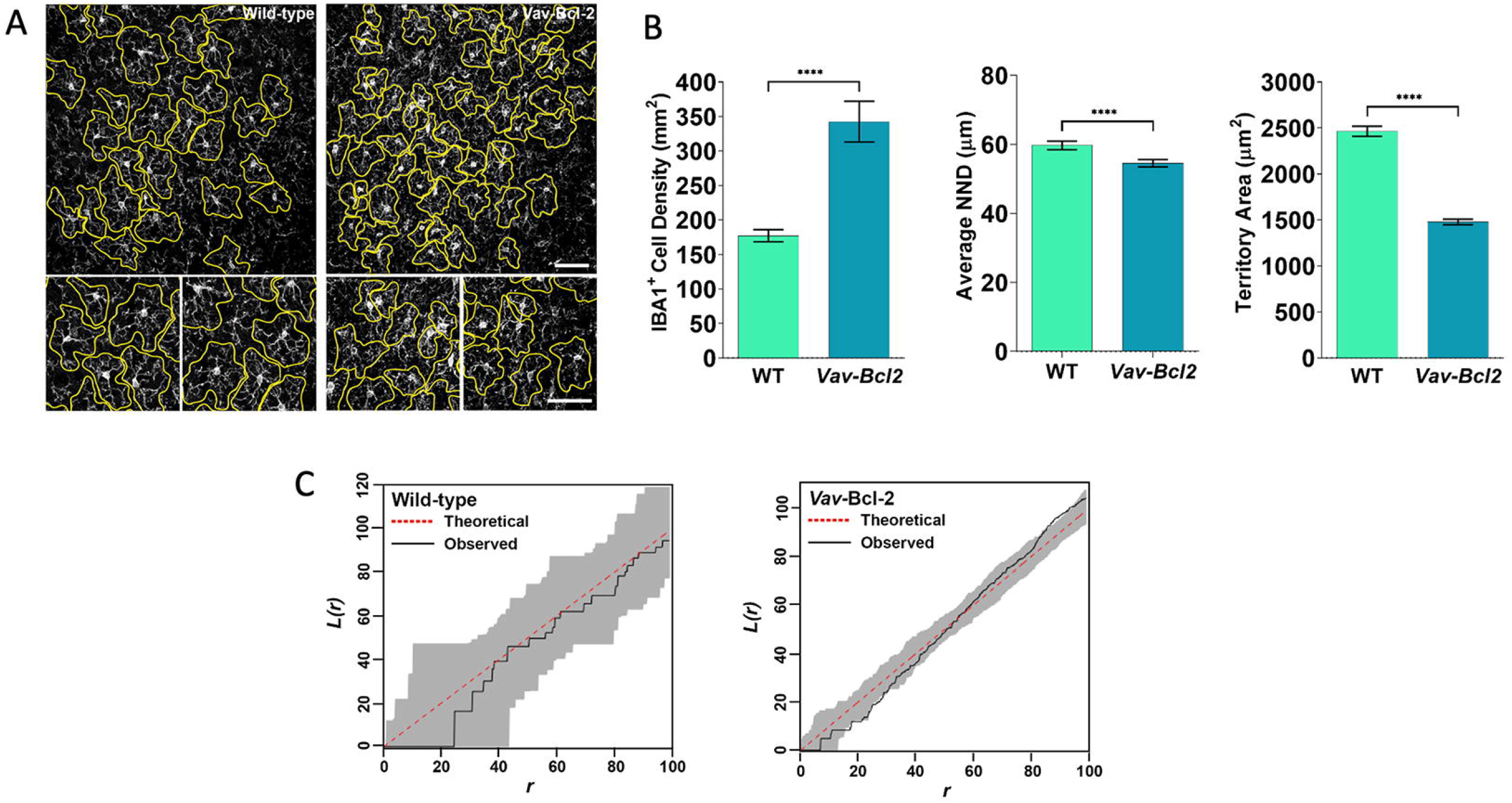
Impact of apoptosis-deficiency on the establishment and maintenance of the microglial mosaic distribution. (A) Microglial territories were traced and measured in 30μm z-stacks of Iba1-labelled brain sections from adult (4-6 months) Vav-Bcl-2 mice (n=4; 312 individual cells) and wildtype (n=4; 470 individual cells) littermates. (B) Quantification of microglial density (Iba1+ cells) in the cortex of adult Vav-Bcl-2 mice and wild-type littermates. Quantification of microglial territory area (μm) in the cortex of Vav-Bcl-2 mice and wild-type littermates. Quantification of the regularity index of microglia in the cortex of Vav-Bcl-2 mice and wild-type littermates. Data shown is represented as mean±SEM. Data were analysed with a student’s t test. Statistical differences: ****p<0.0005. Scale bar in (A) 100μm. (C) Representative plots of the L(r) derivative of Ripley’s K statistics analysing the spatial clustering of microglia from wild-type and Vav-Bcl-2 mice.

### Sparsely labelled microglia expand symmetrically as clonal clusters from early embryogenesis

Our previous data on cell clustering indicated that the dynamics of the microglial population in development could be driven by local clonal expansion events. Therefore, we devised a protocol to achieve very sparse fate-mapping of the embryonic microglial pool with an EGFP label using the *Cx3cr1*^*CreER*^; *Rosa*^*mTmG*^ mouse model. We induced recombination with a low dose of tamoxifen (12.5 mg/kg) to achieve a sparse level of GFP expression in CX3CR1^+^ cells in subsequent offspring, as indicated by the initial optimisation (Supplementary Figure 3). Subsequent analysis of sparsely labelled microglia within the brain primordium was carried out at E11.5, E16.5 and P30 (Figure 3A). GFP^+^ microglia were primarily found as regionally isolated singlets throughout the brain at E11.5 (24 hours post-injection; hpi) with an estimated recombination rate of 6%, indicating that only a small percentage of microglia was initially tagged (Figure 3B,C). Spatial analysis at E11.5 showed that GFP^+^ microglia were dispersed with a very high NND between cells (Figure 3D,E). Expansion of these sparse-labelled microglia was evident at E16.5 and P30 as indicated by a significant increase in the density of GFP^+^ microglia in all brain regions (forebrain, midbrain and hindbrain) (Figure 3C). In spite of the observed increase in density, the recombination rate remained relatively stable at 6.4% at E16.5 and 13.2% at P30 with no significant differences between time points (Supplementary Figure 4).

**Figure 3.**
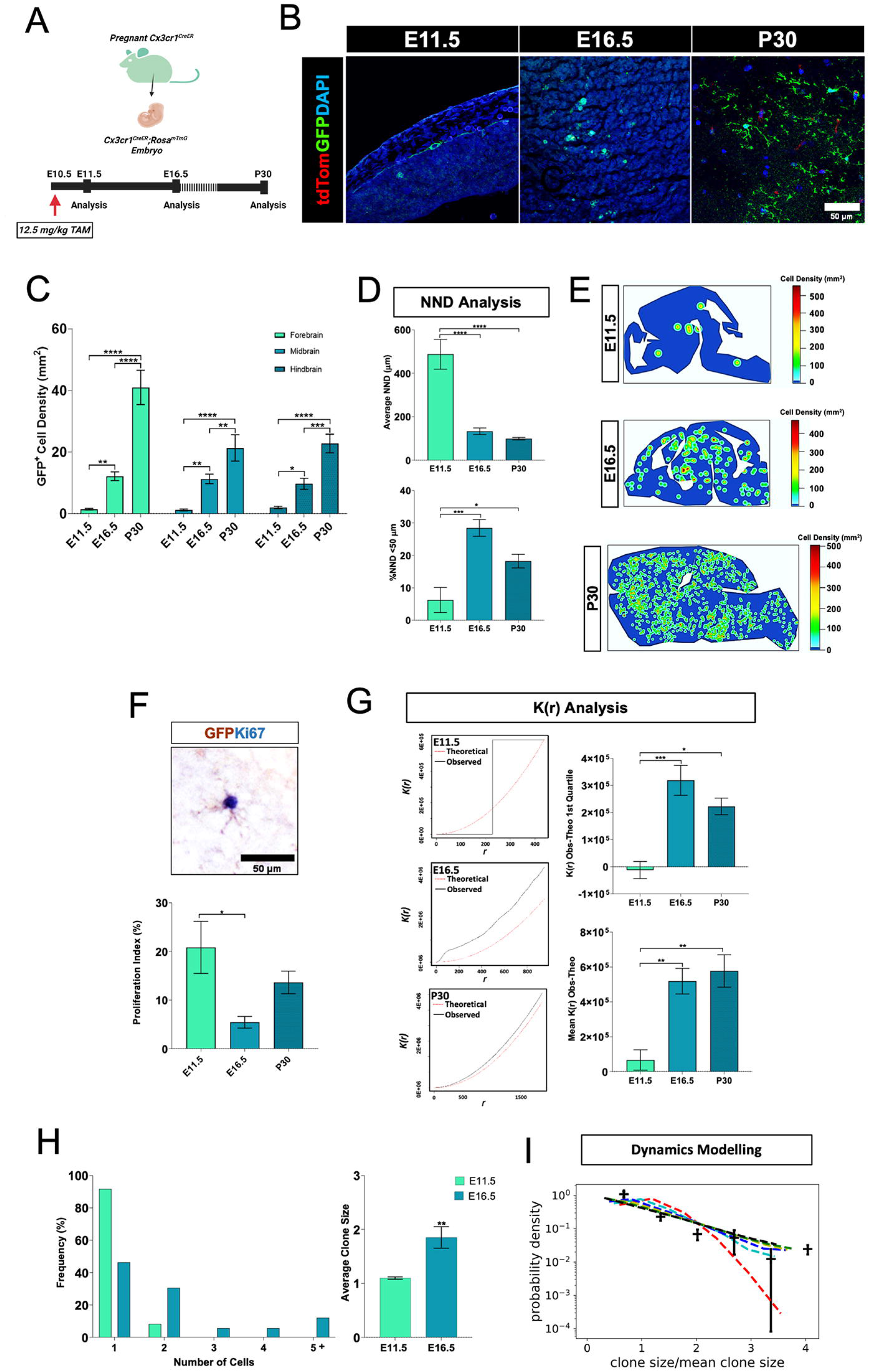
Sparse fate mapping of early embryonic microglia. (A) Experimental overview: induction of recombination in double transgenic Cx3cr1^CreER^; Rosa^mTmG^ embryos by pulsing pregnant dams with either with 62.5 mg/kg of TAM at E9.5 or 12.5 mg/kg of TAM at E10.5 followed by analysis 24 hours post injection (hpi). (B) Representative images of CX3CR1^+^ cells stained with anti-GFP (green) in following the induction of sparse labelling at E10.5 in Cx3cr1^CreER^; Rosa^mTmG^ mice at E11.5, E16.5 and P30. (C) Time-course analysis of the density (cells/mm^2^) of GFP^+^ cells in the forebrain, midbrain and hindbrain at E11.5, E16.5 and P30. (D) Analysis of the NND and %NND<50μm of GFP^+^ cells at E11.5, E16.5 and P30. (H) Representative density maps of GFP^+^ cells at E11.5, E16.5 and P30. (E) Representative image of a GFP^+^PU.1^+^ cell at P30 and analysis of the recombination rate (%GFP^+^cell density/PU.1^+^ cell density). (F) Representative image of a GFP^+^Ki67^+^ cell at P30 and analysis of the proliferation index (%GFP^+^Ki67^+^ cell density/GFP^+^ cell density). (G) Representative K(r) functions of GFP^+^ cells at E11.5, E16.5 and P30. Normalised Ripley’s K analysis of GFP^+^ cells over the entire K(r) function and over the first quartile. (H) The average clone size and clone size frequency was quantified at E11.5 and E16.5. (I) Mathematical modelling of clone size distribution at E16.5 whereby points are probability densities of measured clone sizes and error bars are standard deviations according to standard count stochasticity (Poissonian approximation). The horizontal axis shows the clone size *n* normalised by the mean clone size 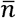, that is,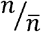. Dashed lines are predictions from model for different values of symmetric division propensity r. Red: r = 0, Cyan: r = 0.2, Blue: r = 0.4, Yellow: r = 0.6, Orange: r = 0.8, Black: r = 1.0, other parameters: *λ* = 0.4 *day*^−1,^ *d* = 0, *δ* = 1; (values of *δ, d* are chosen for simplicity, since the asymptotic clone size distribution for a branching process, as this model is, does not depend on those values(Parigini and Greulich, 2020a). *λ* is estimated from fitting the growth curve of total cell numbers during embryonic development in the forebrain, Fig. 1C). Scale bars are 50 μm and 1000 μm in (B), (E) and (F). Data are expressed as mean ± SEM, E11.5=5, E16.5= 8, P30=5, Two-way ANOVA followed by Tukey’s post-hoc analysis (C) or One-way ANOVA followed by Bonferroni’s post-hoc analysis (E, F, G) or student’s t-test for independent means (H). *p<0.05, **p<0.01, ***p<0.001, ****p<0.0001. Scale bars are 50 μm.

Interestingly we observed that the expansion of the GFP^+^ microglial population was associated with a change in their spatial distribution as GFP^+^ microglia were found in tight clusters at E16.5 suggesting a clonal expansion of the originally labelled isolated progenitors (Figure 3D, 3E). This change in spatial distribution was reflected by changes in the average NND and K(r) of microglia which decreased and increased respectively indicating a clustered distribution (Figure 3D, 3G). Surprisingly, we observed very large ‘clouds’ of GFP^+^ microglia at P30 which spanned entire anatomical areas (Figure 3E). This was unexpected considering the low level of initial labelling at E11.5. Although this technique did not allow us to investigate this further, due to the likely spatial overlap of large clones, it is likely that there is a substantial level of migration to facilitate the widespread distribution of cells observed at P30. Moreover, these data also suggest that early embryonic microglia progenitors are highly proliferative generating large numbers of daughter cells which are capable of colonising the entire brain in a manner that is not restricted to anatomical compartments. In support of this, we observed that the proliferative index of GFP^+^ microglia was relatively high throughout development (Figure 3F). Further spatial analysis at P30 revealed that GFP^+^ microglia continue to display a clustered distribution as indicated by a positive average K(r) value. However, we observed a non-significant decrease of the %NND <50 μm and corrected K(r) over the first quartile which indicates that the GFP^+^ labelled cells are not found as close together and shift towards a more random distribution compared to E16.5 (Figure 3G).

Clonal analysis of sparsely labelled microglia was carried out at E11.5 and E16.5 under the assumption that spatially related GFP^+^ cells derived from the same progenitor. The average size of GFP^+^ clones increased significantly by E16.5 with a range of 1 to 7 cells per clone observed at E16.5 (Figure 3H). To gain insight into the fate of microglia, we applied the Markov model to the clonal data at E11.5 and E16.5 in order to test whether microglia are expanding symmetrically or asymmetrically, i.e. if newly born daughter cells continue to proliferate or whether they exit the cell cycle. Here, the clone-size distribution data at E16.5 fit a model whereby at least a portion of the population undergoes symmetric cell division (Figure 3I) which agrees with our findings that the microglial population expands exponentially during embryonic development. It should be noted that this model does not rule out the contribution of asymmetric divisions. However, the clonal data did not fit a model consisting solely of asymmetric divisions.

### Multicolour fate mapping reveals clonal expansion of postnatal microglia with a disparity in clone size

Our previous results suggested clonal expansion as a primary driver of the dynamics of the microglial population in development. To test this hypothesis more directly, we took advantage of RGB marking, a multicolour labelling strategy, which allows fate mapping of individual cells and their progeny. We first used SFFV-RGB vectors encoding the fluorescent proteins (FPs) mCherry (red), Venus (green) and EBFP2 (blue) under the expression of the ubiquitous spleen form focus virus (SFFV) promoter. Due to the ubiquitous activity of the SFFV promoter, labelling is also expected in non-microglial populations (Gomez-Nicola et al., 2014).

For postnatal fate-mapping of the microglial population, the SFFV-RGB vectors were stereotactically injected into the lateral ventricle at P0 and analysed at P21. Here, RGB labelled microglia were evident in the parenchyma with labelled cells sharing the same RGB colour hue indicating clonal expansion of microglia from common parental cells (Figure 4A). The labelled microglia could be identified due to their unique ramified morphology and co-expression of IBA1 (Supplementary Figure 5). Moreover, an exceptionally diverse colour palette was observed (Supplementary Figure 6) with an RGB colour dispersion and distribution in line with *in vitro* transduction of N13 cells with SFFV-RGB vectors (Supplementary Figure 7) and previous work where RGB lentiviral vectors have been used for clonal analysis (Weber et al., 2011). To define clonal groups, we devised an *in vitro* assay using the N13 cell line. Briefly, SFFV-RGB labelled microglia were seeded at very low densities (100-1000 cells/well) to induce the growth of clonal colonies. The colour spread (AU) in the RGB colour space was calculated for each cluster. Overall, we observed that single-coloured clusters had a low colour spread (<11 AU) indicating that analysis of the RGB colour space was sufficient for clonal discrimination (Supplementary Figure 7). Therefore, we carried out a clonal analysis of labelled microglia at P21 where clones were defined as a group of microglial cells sharing the same colour hue. Our analysis revealed that microglia give rise to clones of heterogenous sizes with the majority of clones consisting of between 1 and 5 cells (Figure 4B). In contrast, a smaller proportion of the clones consisted of much larger numbers of microglia of up to 28 cells, suggesting that there is a disparity in the proliferative index of labelled microglial cells, with some postnatal microglia expanding at a much greater rate than others, and a portion of the population not engaging in proliferation in the postnatal brain (Figure 4B).

**Figure 4.**
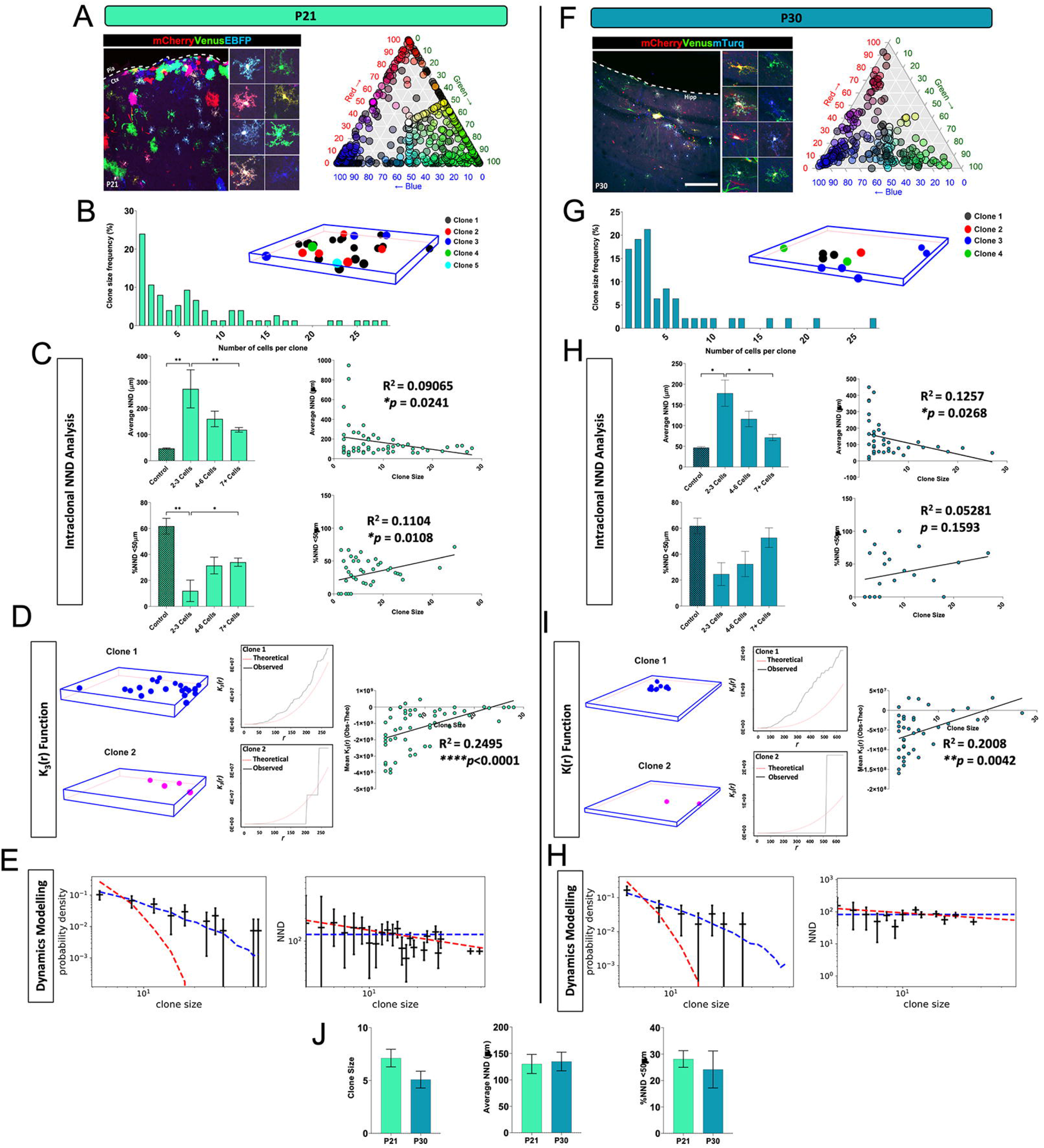
The clonal dynamics of postnatal microglia. (A) Representative example of SFFV-RGB labelled cells in the cortex at P21 including ramified microglia which display a range of multicolour labels (right panel). (B) Quantification of clone size frequency (%) of microglial clones at P21 (n=618 cells) and representative 3-D point pattern of SFFV-RGB labelled microglia where each cell is represented by a different coloured sphere according to the assigned clone it belongs. (C) Quantification of the average intraclonal NND (µm) and %NND <50 µm versus clone size at P21 (n=56 clones). (D) Representative example of a 3-D point pattern and corresponding K(r) function for a large microglial clone (upper) and a smaller microglial clone (lower) with quantification of the mean K(r) (Obs-Theo) versus clone size (n=56 clones) at P21. (E) Mathematical modelling of the fate dynamics of microglia based on their clone size distribution at P21 and NND of clones with the dashed line representing predictions of models of models 1 (red), 2 and 3 (blue). Parameters are chosen arbitrarily as *λ* = 1 *day*^−1^, *r* = 0.5, *δ* =1, *d* = 0 since the model predictions for model versions 2 and 3 are independent of parameters for sufficiently large times (Klein et al., 2010). (F) Representative example of mirRGB labelled cells in the cortex at P30 including ramified microglia which display a range of multicolour labels (right panel). (G) Quantification of clone size frequency (%) of microglial clones at P30 (n=248 cells) and representative 3-D point pattern of mirRGB labelled microglia where each cell is represented by a different coloured sphere according to the assigned clone it belongs. (H) Quantification of the average intraclonal NND (µm) and %NND <50 µm versus clone size at P30 (n=39 clones). (I) Representative example of a 3-D point pattern and corresponding K(r) function for a large microglial clone (upper) and a smaller microglial clone (lower) with quantification of the mean K(r) (Obs-Theo) versus clone size (n=39 clones) at P30. (J) Mathematical modelling of the fate dynamics of microglia based on their clone size distribution at P30 and NND of clones with the dashed line representing predictions of models of models 1 (red), 2 and 3 (blue). (K) Comparison of the average microglial clone size, average NND (μm) and %NND <50 µm between P21 (n=76 clones from 3 animals) and P30 (n=49 Clones from 5 animals). Data are expressed as mean ± SEM, One-way ANOVA followed by Bonferroni’s post-hoc analysis (C and H) or Linear regression analysis and Pearson’s correlation test (C and H) or student’s t-test for independent means (J). Statistical differences: *p<0.05, **p<0.01, ****p<0.0001.

To validate and refine our approach, we next used mirRGB vectors expressing either mCherry (red), Venus (green) or mTurquoise (blue) under the phosphoglycerate kinase (PGK) promoter, and showing superior microglial specificity (in the adult brain 81.26 ± 14.15% of n=228 RGB-positive cells in 4 mice were P2Y12-positive microglia; data not shown).The vectors were stereotactically injected into the lateral ventricle at P0 and analysed at P30. RGB labelled microglia were detected in the brain (Supplementary Figure 8) and similarly to P21, clonal expansion of the RGB labelled microglia could be seen (Figure 4F). We noted that the mirRGB labelled microglia at P30 were reminiscent of SFFV-RGB labelled microglia, including the diverse array of RGB combinations among labelled microglia (Figure 4A,F). Another feature of mirRGB clones that mirrored SFFV-RGB clones was the heterogeneity in clone size with the majority of clones consisting of <5 cells and a minority containing 15+ cells echoing the earlier finding that the proliferation rate of clones is not equal (Figure 4G). Altogether we found no significant differences between the average clone size at P21 (7 cells) and P30 (5 cells) (Figure 4K), indicating that the clonal dynamics of microglia remain stable during this window which is in line with the reported stabilisation of microglial numbers by the third postnatal week (Askew et al., 2017; Nikodemova et al., 2015).

### The microglial mosaic is made up of a diverse patchwork of microglial clones of varying proliferative potential

Next, we took advantage of our RGB marking approach to look more closely at a clonal level and thoroughly investigate the intricate cell-to-cell makeup of the microglial mosaic. At both P21 and P30, we observed a diverse array of spatial patterns among clones, with some appearing to be spatially organised according to clonal groups and others displaying ‘clonal mixing’ with cells of different clones interlocking with others. To objectively characterise the spatial organisation of microglial clones, we calculated the average NND between cells of individual clones and compared it to the average NND of adult microglia (naïve). Interestingly we found that the average NND of clonal microglia at both P21 (130 μm) and P30 (134 μm) is higher than that of naive microglia (46 μm) (Figure 3C and 3H). Moreover, we observed no significant differences between the average NND distance of clonal microglia at P21 or P30 suggesting that the spatial distribution of clones is stable during the juvenile phase (Figure 4K). A similar trend was seen when we assessed the % of microglia with a NND of less than 50 μm in both clonal microglia and naïve microglia, with 61% of naïve microglia displaying a <NND 50μm compared to 28% (P21) and 24% (P30) in clonal microglia (Figure 4K).

Spatial analysis revealed a significant negative correlation between clone size and the average NND of clones at both P21 and P30, indicating that microglial cells from larger clones typically have a smaller NND in comparison to those of smaller-sized clones which are further apart (Figure 4C,H). Similarly, we found a positive trend between clone size and %NND <50μm which reached statistical significance at P21 (Figure 4C,H). This finding indicates that microglia from larger clones display a spatial association which we were able to confirm by applying the spatial statistics test, Ripley’s K (K(r)) function, to analyse our 3D stacks (Figure 4D,I). Here we found that larger clones were more clustered compared to smaller clones which were typically dispersed as evidenced by a positive trend between clone size and the mean K(r) and normalised K(r) (Observed-Theoretical) which reached significance at P21 (Figure 4D,I). In sum, we did not find any evidence of ordered organisation of microglial clones. However, our spatial analyses suggest that larger clones dominate spatial niches within the microglial mosaic in comparison to smaller clones which are dispersed and ‘diluted’ throughout the network.

To gain insight into the proliferative kinetics of clonal microglia, we applied mathematical models to the clonal data at P21 and P30. We initially considered three models: 1) Microglia are actively dividing all at the same time and clones are equipotent; 2) microglia are initially quiescent and start dividing at random times with equipotent clones; and 3) microglia start dividing at the same time but with different proliferative activity ν, which is inherited along with a clone, i.e. some clones have a sustained proliferative advantage over others. For our modelling, we assumed that cell division rates are uniformly distributed. We observed that model variant (1) does not fit the clonal data, but both model (2) and model (3) do (notably the model outcomes are parameter-free when we use consider probability densities and normalised clone sizes) (Figure 4E,J). Hence, we can exclude that cells start dividing at the same time and are equipotent. The observation that both models (2) and (3) have the same output is reasonable, since a shorter period of divisions is dynamically equivalent to the same time period but a larger division rate among all cells in a clone. To distinguish the latter models, we utilised an analytical model based on the NND data of cells within clones. We next considered two scenarios: 1) Clones are equipotent, that is, on average, no clone has a proliferative advantage (while individual cells may divide faster, any proliferative advantage is not inherited). Clones may initiate proliferation at any time (no distinction between model variants (1) and (2) above); 2) clones are heterogeneous, i.e. the proliferative potential, and thus ν differs between clones. Here, we found that models do not significantly vary from the data, however, model (2) produces overall a better fit for time point P21 suggesting that the proliferative potential is heterogenous among microglial clones (Figure 4E,J).

## DISCUSSION

Understanding the proliferative mechanisms employed by the microglial population in both development and disease has garnered a lot of interest in recent years (Gomez-Nicola et al., 2013; Fuger et al., 2017; Tay et al., 2017b; Hammond et al., 2021). In the healthy brain, resident microglia self-renew in a stochastic manner throughout life, with varying rates of turnover in different brain regions (Lawson et al., 1990; Askew et al., 2017; Fuger et al., 2017; Tay et al., 2017a). In response to any deviation from homeostasis such as in disease (Tay et al., 2017a; Jordão et al., 2019) and in depopulation-repopulation paradigms (Bruttger et al., 2015; Zhan et al., 2019), microglia increase their rate of proliferation and may undergo clonal expansion in order to increase their cell numbers. Here, we expand our understanding of the proliferative dynamics of microglia during development and demonstrate for the first time that developmental growth of the microglial population is facilitated by clonal expansion of early precursors during embryonic and postnatal developments which form an intricate patchwork of clones within the microglial mosaic. Moreover, the general expansion of microglia can be described as allometric, whereby the increase in microglial numbers is proportional to the increasing size of the brain.

The number of studies investigating clonal dynamics in microglia is limited, mainly due to technological limitations associated with multicolour reporter lines, seen for example in the Confetti strain (Tay et al., 2017; Zhan et al., 2019). While such studies have been informative, the Confetti model for example is limited to 4 colour combinations and therefore lacks the resolution to provide an in-depth clonal analysis. Here we used a combination of SFFV- and MIR-RGB vectors to label microglia in the postnatal brain which resulted in a multitude of colour combinations and thus allowed us to thoroughly characterise the clonal dynamics of these cells. Our subsequent clonal analysis unveiled heterogeneity in clone size, with a small subset of microglia generating very large clones. This suggests a considerable degree of clonal variability linked to the proliferative potential: final clonal size will be instructed by the proliferation capacity of the originally labelled parent cells, and then their progeny. Interestingly, similar proliferative heterogeneity has also been reported in a model of microglial depletion/repopulation in the visual cortex (Mendes et al., 2021). In the context of the existing literature, our findings raise a number of questions including the source of this proliferative diversity and whether or not this is a long-term feature of individual microglial cells that is carried through until adulthood.

We also utilised a method of sparse-labelling to track the progeny of individual microglia from early embryogenesis. Our findings indicate that some individual microglia undergo mass expansion as evidenced by the clouds of sparse-labelled microglia at P30. This finding is in line with a recent study where *in vivo* barcoding was used to track clonal expansion of various cell types, including microglia, within the embryonic brain (Ratz et al., 2022). In line with our findings, this study showed that expanded microglial progenitors span anatomical niches in an almost continuum (Ratz et al., 2022). Here we examined the changes in the spatial distribution of sparsely labelled microglia, which were highly clustered during embryonic development in comparison to adulthood, when the labelled microglia were more randomly distributed with a lower NND in line with the formation of the mosaic similar to the trends observed in the total PU.1^+^ population. The drivers of these changes in spatial distribution remain unclear, although our data and those of others suggest that the attraction and repulsion of microglia to either dying or proliferating cells may facilitate microglial dispersion during development (Casano et al., 2016; Cunningham et al., 2013; Hattori et al., 2020). On this note, we observed that RGB marked clones displayed a level of spatial mixing indicating that the microglial mosaic is not made up of strict clonal units. However, spatial analysis of RGB clones revealed that larger clones tended to be closer together in space and begs the question of whether there is a level of clonal dominance in certain niches throughout the brain. This may be important if functional mutations are inherited by daughter cells of a clone which may render certain spatial niches vulnerable to microglial dysfunction.

An interesting observation from this study was the apparent relationship between microglial expansion and space availability. Our results suggest that there is a coupling between the rate of microglial expansion and the growth rate of the brain resulting in a plateaued cellular density during embryonic development. A similar plateau was reported in a recent study where the changes in microglial density were investigated in the hypothalamus (Marsters et al., 2020). The mathematical modelling applied here supports that during embryonic development there is an exponential growth phase, during which microglia primarily undergo symmetric divisions similar to what has been observed during embryogenesis in the zebrafish (Svahn et al., 2013). In contrast to embryonic development, we observed a remarkable increase in the number and density of microglia during postnatal life with a peak in microglial density at P14, in line with previous reports (Alliot et al., 1999; Nikodemova et al., 2015; Wlodarczyk et al., 2017). Interestingly, the increase in cell number still tracks a trend of increasing brain area. This type of growth in a cell population is referred to as allometric growth and is reported in a variety of cell types across species and is important for regulating the size of certain organs (Vollmer et al., 2017). The phenomenon of allometric growth is not thoroughly understood and is thought to be influenced by both intrinsic and extracellular mechanisms of growth control (Vollmer et al., 2017). The intricate connection of niche availability and stemness potential has recently been defined as a mechanism of ‘crowding feedback’, whereby under suitable feedback regulation, the property of ‘stemness’ is entirely determined by the cell environment (Greulich et al., 2021).

Indeed, in further support of this hypothesis, we found that changes in the density of microglia correlated with changes in the NND of cells. This suggests that the density of microglia will increase during development until cells become associated with each other, in other words when there is an equilibrium, and the mosaic network is achieved. This observation sheds light on the reported ability of microglia to recapitulate their normal densities and spatiotemporal distribution following depletion in several paradigms (Elmore et al., 2014; Bruttger et al., 2015; Zhan et al., 2019). Considering the interplay between brain growth and microglial expansion, it is not unreasonable to predict that space availability may favour the expansion of certain microglial progenitors over others that are spatially constricted and thus may explain the observed disparity in clone size at P21 and P30.

Furthermore, these findings suggest contact inhibition as an underlying mechanism. Contact inhibition is a process of allometric growth control that has been described in cell culture and cancer cells, whereby cells will expand and migrate freely until they come into contact with another cell or physical barrier (Mendonsa et al., 2018; Pavel et al., 2018; Tsuboi et al., 2018). The mechanisms of contact inhibition are thought to rely on membrane-bound cell adhesion molecules (CAMs) such as E-cadherin. For example, loss of E-cadherin disrupts contact inhibition resulting in metastasis of epidermal cells (Navarro et al., 1991). Currently, there are limited data regarding potential pathways regulating contact inhibition in microglia and it is unclear whether CAMs are involved in this. One potential transmembrane proteoglycan which may be involved is Syndecan-4 which has been previously implicated in regulating contact inhibition in mesenchymal cells (Valdivia et al., 2020). Interestingly, it was observed that Syndecan-4 is downregulated in microglia following their depletion which led the authors to suggest a potential role for Syndecan-4 in regulating microglial contact inhibition (Zhan et al., 2019). Other potential sources of allometric growth control are spatial checkpoints in the cell cycle such as those reported between the G1 and S phase in epithelial cells which will prevent proliferation when cells become spatially restricted (Streichan et al., 2014).

Taken together, our findings demonstrate that the developmental expansion of microglia is driven by clonal expansion of early invading microglial progenitors, with a small proportion generating relatively large clones. The density of microglia rapidly increases during postnatal development until a relatively even spatial distribution: the microglial mosaic is achieved. In addition, spatial analysis reveals that the microglial mosaic is fashioned from an intricate patchwork of diverse microglial clones. These findings uncover another layer of the proliferative capabilities of microglia and thus will be useful for understanding the behaviour of these cells during neurodevelopmental and neurological diseases.

## ACKNOWLEDGEMENTS

We thank the Southampton Flow Cytometry Facility and the Imaging Unit for technical advice and the Biomedical Research Facility for assistance with animal breeding and maintenance. We thank Georgina Dawes for technical assistance; Maria Olmedillas del Moral, Cris Richter and Bianca Brawek for analyses of the specificity of MIR-RGB vectors. We thank Professor Mark Cragg for provision of vav-Bcl2 mice. The research was funded by the Leverhulme Trust (RPG-2016-311) and the Medical Research Council (MRC) (MR/P024572/1).

## AUTHOR CONTRIBUTIONS

DG-N designed the study, secured the funding and supervised the project. LB-C performed most of experiments and analysed the data. RM and KEA contributed experimental work and analysis. KR provided SFFV-RGB vectors. K.L and O.G. provided the MIR-RGB vectors. PG performed mathematical modelling of the data. LB-C, DAM and DG-N wrote the manuscript. All authors contributed to drafting the manuscript.

## FIGURE LEGENDS

**Supplementary Figure 1**.

(A) Immunostaining of sections from E16.5 embryos was carried out with anti-PU1 and anti-IBA1 as markers of microglia as well as with anti-Caspase3, anti-Ki67 and anti-Olig2 as markers of dying cells, proliferating cells and OPCs respectively. (B) Representative point patterns were generated in order to visualise the localisation of microglia with Caspase3^+^, Ki67^+^ and Olig2^+^ cells. N=3; scale bar is 100 μm.

**Supplementary Figure 2**.

(A) Linear regression analysis between the density and average NND (μm) of microglial cells during development. (B) Linear regression analysis between the density and %NND <50 μm of microglial cells during development. N=3 per group, Linear regression analysis and Pearson’s correlation test. Statistical differences: *p<0.05.

**Supplementary Figure 3**.

(A) Experimental design. (B) Representative images of GFP^+^ (brown) and PU.1^+^ (blue) cells. (C) Comparison of the recombination rate between both setups. Data are expressed as mean ± SEM, E9.5=4, E10.5=5, student’s t-test for independent means. Each timepoint consists of a single litter. Statistical differences: **p<0.01. Scale bar is 50 µm.

**Supplementary Figure 4**.

(A) Representative images of GFP^+^ (brown) and PU.1^+^ (blue) cells at P30. (B) Comparison of the recombination rate between E11.5, E16.5 and P30. Data are expressed as mean ± SEM, E11.5=5, E16.5= 6, P30=5, One-way ANOVA followed by Tukey’s post-hoc analysis. Each timepoint consists of a single litter. Scale bar is 50 μm.

**Supplementary Figure 5**.

Representative example of IBA1^+^ cells (cyan) successfully transfected with the SFFV-RGB vectors encoding the fluorescent proteins mCherry (red), Venus (green) and EBFP (blue) at P21. Scale bar is 100 μm.

**Supplementary Figure 6**.

(A) Quantification of the colour intensity (%) in the red, green and blue channels from all SFFV-RGB labelled microglia at P21 (N=618 cells from 3 animals). (B) The overall proportion of RGB labelled microglia that were single, double or triple labelled with SFFV-RGB vectors. (C) Quantification of the colour intensity (%) in the red, green and blue channels from all MIR-RGB labelled microglia at P30 (N=249 cells from 5 animals). (D) The overall proportion of RGB labelled microglia that were double or triple labelled with MIR-RGB vectors. One-way ANOVA followed by Tukey’s post-hoc analysis. Statistical differences: ***p<0.001, ****p<0.0001.

**Supplementary Figure 7**.

(A) Experimental design. (B) Representative images of N13 cells transduced with SFFV-RGB vectors encoding the fluorescent proteins mCherry (red), Venus (green) and EBFP (blue). (C) The corresponding colour space of RGB labelled N13 cells. (D) Quantification of the colour intensity (%) in the red, green and blue channels from all SFFV-RGB N13 cells. (E) The overall proportion of RGB labelled N13 cells that were double or triple labelled with SFFV-RGB vectors. N=615 cells; One-way ANOVA followed by Tukey’s post-hoc analysis. Statistical differences: ****p<0.0001. Scale bar is 100 μm.

**Supplementary Figure 8**.

Representative example of IBA1^+^ cells (cyan) successfully transfected with the MIR-RGB vectors encoding the fluorescent proteins mCherry (red), Venus (green) and mTurq (blue) at P30. Scale bar is 50 μm.

## METHODS

### Mice

All mice were maintained according to Home Office regulations and experiments were approved by a local ethical review committee in accordance with the UK animals (Scientific Procedures) Act (1986), and conducted under relevant personal and project licenses. All mice used were bred on a C57BL/6 background. C57BL/6 and *c-fms* EGFP mice (Sasmono and Williams, 2012) (hereafter referred to as Macgreen) were used for time course analysis of microglia throughout embryonic and postnatal life. For lineage tracing experiments, homozygous Rosa^mT/mG^ (Muzumdar et al., 2007) males were crossed with homozygous Cx3cr1^CreER^ (Goldmann et al., 2013) female mice to generate Rosa^mT/mG^ x Cx3cr1^CreER^ mice (hereafter referred to as Cx3cr1^CreER^; Rosa^mT/mG^). *Vav-Bcl2* mice (Egle et al., 2004) were used as a model of myeloid-specific apoptosis prevention, as previously described for microglia (Askew et al., 2017). C57BL/6 mice were used for all *in vivo* experiments involving stereotaxic injection of vectors for RGB marking.

For developmentally-timed experiments, trios were set up late in the afternoon. Following this, female mice were checked daily for a vaginal plug. The day on which the plug was identified was noted as embryonic day 0.5, or E0.5, estimating the time of copulation to be 12 hours after initial pairing based on the light-dark cycle and mating habits of mice (Behringer et al., 2014). Female mice that had a vagina plug were separated from males.

### Cell lines

The N13 cell line of microglial cells was used for the characterisation of RGB marking using LeGO vectors (Gomez-Nicola et al., 2014; Weber et al., 2012; Weber et al., 2011). The cells were cultured according to manufacturer’s guidelines in Dulbecco’s Modified Eagle’s Medium (D-MEM) that was supplemented with 10% fetal bovine serum (FBS) and Penicillin-Streptomycin (all ThermoFisher Scientific).

### Induction of cre-dependent sparse labelling of microglia with EGFP

Cx3cr1^CreER^; Rosa^mT/mG^ mice were used for sparsely labelling microglia with an EGFP tag. The Rosa^mT/mG^ transgene contains the fluorescent proteins tdTomato and EGFP under the Rosa26 promoter (Goldmann et al., 2013; Muzumdar et al., 2007). For sparse labelling microglia with EGFP during development, we first prepared stocks of 20 mg/ml and 10 mg/ml of TAM and 4-OH TAM by solubilising in ethanol (100%) and dissolving in corn oil (Sigma-Aldrich) (1 hr; 60°C). TAM and 4-OH TAM were sonicated using a bench top vortex until completely solubilised. The stock solution of TAM was supplemented with 10 mg of progesterone to offset the reported off-target effects of TAM during pregnancy (O’Koren et al., 2019). Using low doses of TAM allowed labelling of a very small percentage of microglia arising during early embryonic development. Pregnant Cx3cr1^CreER^ dams that had been mated with male Rosa^mT/mG^ mice received an i.p. injection of TAM (200 mg/kg, 62.5 mg/kg or 12.5 mg/kg) on either E9.5 or E10.5. Dams were weighed and closely monitored following injection for any signs of sickness behaviour or resorption of pregnancy. For optimisation purposes, the subsequent offspring were analysed 24 hours post injection (hpi) in order to estimate the initial rate of EGFP labelling in CX3CR1^+^ cells. Following optimisation, a time course was established following i.p. injection of TAM (12.5 mg/kg) to pregnant dams at E10.5 with subsequent analysis of sparse labelled microglia in Cx3cr1^CreER^; Rosa^mT/mG^ offspring at E11.5, E16.5 and P30.

### Multicolour fate mapping with RGB vectors

#### Production and use of VSVG-SFFV-RGB vectors

RGB marking of cells can be achieved by transduction with a combination of LeGO vectors encoding a red, green and blue fluorescent protein (Weber et al., 2011) hereby referred to as RGB vectors. LeGO vectors are 3^rd^ generation, self-inactivating lentiviral vectors derived from HIV-1, containing all necessary cis-active elements for packaging, reverse transcription and integration into the host cell genome and are therefore suitable for the stable and long-term labelling of mammalian cells with fluorescent proteins (Weber et al., 2011). Due to the random nature of the transduction with the tree RGB vectors, target cells display a unique colour hue resulting from a stochastic mixture of the three basic colours, that is inherited by all daughter cells and thus is suitable for clonal analysis (Weber et al., 2011). Moreover, RGB vectors can be engineered for cell specificity by use of cell-specific promoters and can be used for *in vitro* and *in vivo* studies (Gomez-Nicola et al., 2014).

For *in vitro* based RGB marking of N13 cells, SFFV-RGB vectors were used, as the strong and ubiquitous spleen focus forming virus (SFFV) promotor allows stable expression in glial cells (Gomez-Nicola et al., 2014). Packaging with the envelope protein from vesicular stomatitis virus (VSVG) leads to high vector titers and a broad tropism of the viral particles. The titers were further increased by centrifugation at 8,000 g and 4°C overnight. We made available all LeGO vectors used here through Addgene.org: LeGO-EBFP2 (Addgene #85213), LeGO-V2 (Addgene #27340) and LeGO-C2 (Addgene #27339). A combination of SFFV-RGB vectors encoding; enhanced blue fluorescent protein (EBFP2) (titer per ml = 3.39 × 10^9^), Venus (titer per ml = 7.86 × 10^9^) and mCherry (titer per ml =3.39 × 10^9^) were used at an equimolar concentration. N13 cells were initially seeded at low densities (between 10 and 100 cells per well) in order to try and achieve clonal colonies over time. After 24 h, the cells were transduced with 10 µl of this SFFV-RGB vector mix and incubated for a further 24 h. The media was changed in order to remove the RGB vectors and the cells were maintained for 5 more days before fixation and analysis. To analyse the potential colour spread of the RGB-marked cells, a second experimental setup was used where N13 cells were initially transfected with 150 µl of the above SFFV-RGB vector mix while in T75 flasks. The flasks of N13 cells were split 24 h after transfection and subsequently seeded at low densities (1000 or 100 cells per well) in order to induce the growth of clonal colonies. After 6 days, cells were fixed with 2% PFA and washed with PBS before confocal analysis of RGB marked colonies.

#### Production of Lenti-RGB.miR-9.T vectors

For producing the Lenti-RGB.miR-9.T vectors, we modified the parent construct (LV.GFP.miR-9.T) (Åkerblom et al., 2013), which incorporates four target sites (5⍰-TCATACAGCTAGATAACCAAAG-3⍰) for microRNA-9 (miR-9) downstream of GFP, thus causing the degradation of the fluorescent protein-encoding messenger RNA or silencing of the construct’s expression in cells expressing miR-9. Because unlike the other brain cells of neuroectodermal origin (e.g., neurons, astrocytes and oligodendrocytes) microglia lack miR-9 expression, this approach favours selective labelling of microglia (Åkerblom et al. 2013)(Brawek et al., 2017). Lenti-RGB.miR-9.T vectors were generated by replacing GFP with either mCherry, Venus, or mTurquoise2. Cell-free supernatants containing viral particles were produced by transient transfection of HEK293T packaging cells with the lentiviral construct and helper plasmids (psPAX2 and pMD2G) as described previously (Brawek et al. 2017). After 48 h the virus-containing culture supernatant was collected, filtered through a 0.45 μm pore-sized filter and concentrated by centrifugation at 27,000 rpm for 2 h at 4 °C by using Thermofisher WX Ultra80 centrifuge (Waltham, MA, USA). Pellets were re-suspended in sterile PBS and stored at - 80 °C. Viral solutions with a titre higher than 10^8^ colony forming units/ml were used in this study.

#### In vivo transduction of microglia with RGB vectors

SFFV-RGB vectors and Lenti-RGB.miR-9.T vectors (MIR-RGB vectors) were used for *in vivo* tracing of microglia. In order to induce multicolour labelling of microglia during postnatal development, SFFV-RGB vectors encoding the fluorescent proteins mCherry (red), Venus (green) and enhanced blue fluorescent protein (EBFP)(blue) were stereotaxically injected (1 μl) at an equimolar concentration into the right brain hemisphere of postnatal *C57BL/6* pups (n=3) at P0. The injected mice were closely monitored for signs of sickness behaviour and subsequently culled after 21 days for histological analysis. To increase the specificity of transduction in microglia, we used the MIR-RGB vectors under the control of the widely expressed phosphoglycerate kinase (PGK) promoter (Åkerblom et al., 2013). For experiments using MIR-RGB vectors, postnatal *C57BL/6* pups (n=5) received an ipsilateral injection at P0 to the right hemisphere with 1µl of an equimolar mix of MIR-RGB vectors encoding the fluorescent proteins mCherry (red), Venus (green) and mTurquoise (blue). A control group (n=2) did not receive any injection. Following injection, the mice were closely monitored for signs of sickness behaviour. Mice were culled 30 days following injection for histological analysis.

### Immunohistochemistry

Sagittal or coronal brain sections were cut from paraformaldehyde-fixed brains using a vibratome as described previously (Gomez-Nicola et al., 2013). Whole embryos were embedded in paraffin wax and sectioned using a microtome. Sections for brightfield microscopy were initially incubated with Dako dual enzyme block to quench endogenous peroxidase and alkaline phosphatase (AP) activity (Agilent Technologies) (15 min; RT) and then washed with PBST0.1%. Wax embedded sections contained additional dewaxing (60°C oven for 1h and xylene), rehydration (sequential rehydration in ethanol; 100% to dH2O) and antigen retrieval (boiled in citrate buffer for 15 min) steps prior to quenching. To prevent unspecific binding of primary and secondary antibodies, sections were incubated with a blocking solution made in PBST0.2 (0.2% Tween20) and containing 5% bovine serum albumin (BSA) and 5% of the appropriate animal serum of the secondary antibody host animal (1 hr; RT). Sections were incubated with primary antibodies (ON; 4°C) that were appropriately diluted in blocking solution: chicken anti-GFP (Abcam; ab13970; 1:500), rabbit anti-GFP (Santa Cruz Biotechnology; sc-8334; 1:500), rabbit anti-Iba1 (Wako; 1:500), goat anti-Iba1 (Invitrogen; PA518488; 1:500), rabbit anti-Olig2 (Santa Cruz Biotechnology; sc-48817; 1:500), rabbit anti-PU.1 (Cell Signalling Technology; 2258; 1:500), rabbit anti-PU.1 (Santa Cruz Biotechnology; sc-352; 1:500), rabbit anti-Caspase3 (Merck Millipore; PC679; 1:50) or rabbit anti-Ki67 (Abcam; ab15580; 1:500). Sections were washed and then incubated with appropriately diluted biotinylated secondary antibodies (Vector) or Alexa-488, Alexa-568 and Alexa-647 fluorescent secondary antibodies (Invitrogen). For immunofluorescence, sections were incubated with the nuclear counterstain DAPI (1:200) in PB (10 min; RT) and then washed with PB (3 × 5 min; RT) and finally mounted on gelatinised slides and coverslipped with Mowiol DABCO (Sigma). For brightfield microscopy, sections were incubated with the avidin-biotin complex (ABC) kit and visualised with horse-radish peroxidase substrate diaminobenzidine (DAB) (2.5%) and/or BCIP/NBT kit (Vector) according to the manufacturer’s instructions. Methyl green (Vector) was used as a counterstain. Sections were mounted on gelatinized slides and coverslipped with Mowiol DABCO or DPX (Sigma).

### Image acquisition and analysis

Fluorescent signal was detected and visualised using a Leica DM5000 B microscope or a Leica SP8 confocal system. 3-D confocal stacks were acquired from tissues transduced with RGB vectors using the laser scanning function of the Leica SP8 confocal microscope with a 20x 0.75NA objective at a resolution of 1.3 pixels per μm. Z stacks were acquired at 1.14 μm per step in the Z plane from either 35 μm or 80 μm tissue sections. Confocal tile scanning of whole sagittal tissue sections was carried out using the Leica SP8 confocal microscope using the spiral scan function in Leica LAS-X software program. Brightfield images of DAB and AP staining were obtained using a Leica DM5000 B microscope or an Olympus VS110 slide scanner at a resolution of 2.9 pixels per μm.

Images were analysed using Fiji analysis software (Schindelin et al., 2012). The CellCounter plugin was used to record the number of cells and their spatial coordinates (XY). For each antigenic marker, the cell density (cells/mm^2^) was calculated as the overall cell number divided by the area of the ROI (mm^2^) (n=3 sections/mouse). For analysis of double positive cells, the deconvolution tool was used for separation of the chromogenic markers, DAB and AP. Delineated boundaries were saved as an ROI file and the spatial coordinates of each boundary were exported using the ROI manager tool in Fiji.

### Analysis of microglia transduced with RGB vectors

Fiji was used for analysis of RGB fluorescent colour from multicolour labelling experiments in N13 cells and postnatal mice that had been transduced with RGB vectors. An automated macro was developed for analysis of RGB colour from *in vitro* experiments. Briefly, the cell somas of fluorescently labelled microglia were delineated from maximum projection of confocal stacks. For *in vitro* analysis, the colour histogram tool was used to calculate the mean intensity in the red, green and blue channel for microglia cell soma from RGB images. For *in vivo* experiments, a manual analysis was used in order to reduce any noise and avoid artefacts. Images from *in vivo* experiments were converted from composite to RGB images and the channels were split into corresponding red, green and blue images which were displayed in greyscale. The average intensity in the red, green and blue channel for each microglia soma was calculated by measuring the mean grey value for each image. The resultant RGB colour space was mapped onto ternary graphs as a percentage of colour intensity using the Ternary Package in RStudio (Smith, 2017). For *in vitro* analysis of RGB colour spread within clonal clusters, the distance (arbitrary units (AU)) was measured between two of the furthest points, *a* and *b*, on the ternary plot using the distance formula for the *X, Y* and *Z* coordinates of each point:

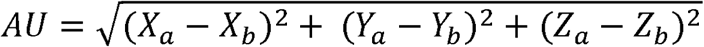

### Spatial analysis using SpatStat

The SpatStat package in RStudio was used for analysis of the spatial distribution of cells (Baddeley et al., 2016). The two-dimensional coordinates (XY) for 2-D images and three-dimensional coordinates (XYZ) for 3-D stacks were saved as .csv files and imported as a dataset into RStudio. Point patterns were generated using the coordinates of each cell and the coordinates delineating the boundary of the area of interest. For spatial analysis of two different cell types from the same image (i.e., PU.1^+^ cells and Caspase3^+^ cells), a subsequent column defining the cell type was included. For analysis of multiple cell types, the ‘mark’ function was used to define the cell type. The point patterns were converted to density heat maps based on a kernel bandwidth of 25 or 50 μm (command ‘Density’ in SpatStat) in order to represent the spatial distribution of cells.

### Nearest Neighbour Analysis

The average nearest neighbour distance was computed using SpatStat (command ‘nndist’) which will compute the nearest distance from a cell to the next in a point pattern. The resultant vector of values was exported as .csv files and used to compute the average nearest neighbour distance. For calculation of the nearest neighbour values from the 1^st^ to 100^th^ neighbouring cell, the range of 1-100^th^ neighbour was specified and returned as the mean distance. The resultant vectors of values were exported as .csv files.

### 2D Ripley’s K spatial analysis

Statistical spatial analysis was carried out using Ripley’s K function (command ‘Kest’ in SpatStat) in order to characterise the spatial distribution of microglia over a given radius. The K function of a stationary point process *X* was defined so that lambda *K(r)* equals the expected number of additional random points within a distance *r* of a typical random point of *X*. Here lambda is the intensity of the process, i.e. the expected number of points of *X* per unit area. The K function was determined by the second order moment properties of *X*. In order to make inferences about the spatial distribution, the estimate of *K* for the observed spatial points was compared to the true value of *K* for a completely random (Poisson) point process, which is:

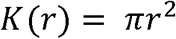

Deviations between the observed and Poisson K curves suggested spatial clustering or spatial regularity. The estimates of K(r) are of the form:

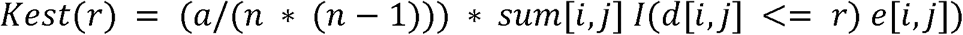

where *a* refers to the area of the window, *n* is the number of data points, and the sum is taken over all ordered pairs of points *i* and *j* in X. Here *d[i,j]* is the distance between the two points, and *I(d[i,j] <= r)* is the indicator that equals 1 if the distance is less than or equal to *r*. The term *e[i,j]* is the edge correction weight which is applied to reduce bias (Ripley, 1988). The correction implemented here is Ripley’s isotropic correction used for rectangular and polygonal windows (Ripley, 1988).

Points were assumed to be randomly distributed when the observed *K(r)* was equal to the theoretical *K(r)*. Points were considered as uniformly distributed or dispersed when the observed *K(r)* was less than the theoretical *K(r)*. Conversely, when the observed *K(r)* was greater than the theoretical *K(r)*, the points were considered clustered. Observed *K(r)* and theoretical *K(r)* values were assessed over the first quartile of each K function. Observed *K(r)* and theoretical *K(r)* values were also calculated as averages over the entire K function. Subtraction of the theoretical *K(r)* from the observed *K(r)* was used for normalisation across different point patterns.

### 3D Ripley’s K spatial analysis

For confocal stacks, Ripley’s K function was estimated from 3-D patterns (command ‘K3est’ in SpatStat). For three-dimensional stationary point processes *Phi*, the three-dimensional *K* function is

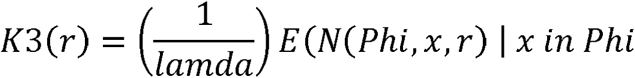

where *lambda* is the intensity of the process (the expected number of points per unit volume) and *N(Phi,x,r)* is the number of points of *Phi*, other than *x* itself, which fall within a distance *r* of *x*. This is the three-dimensional generalisation of Ripley’s *K* function for two-dimensional point processes (Ripley, 1977).

The three-dimensional point pattern X is assumed to be a partial realisation of a stationary point process *Phi*. The distance between each pair of distinct points was computed. The observed cumulative distribution function of these values, with appropriate edge corrections, was renormalised to give the estimate of *K3(r)*. The edge correction implemented here was the three-dimensional counterpart of Ripley’s isotropic correction.

### Marked Connect Function of Multitype Point Patterns

The mark connection function *p[A,B](r)* was used as a measure of the spatial dependence between two cell types of a process *X* at distance *r* apart. The mark connect function *p[A,B](r)* is defined as the conditional probability, given that there is a point of the process at a location *u* and another point of the process at a location *v* separated by a distance *||u-v|| = r*, that the first point is of type *A* and the second point is of type B (Arnold, 1995). If the marks attached to the points of *X* are independent and identically distributed, then *p[A,B](r) = p[A]p[B]* where *p[A]* denotes the probability that a point is of type *A* and *p[B]* denotes the probability that a point is of type *B*. Values larger than this, *p[A,B](r) > p[A]p[B]*, indicate positive association between the two types, while smaller values indicate negative association. For analysis of spatial dependence across animals, the probability of an independent association *p[A]p[B]* was subtracted from the calculated mark connect function *p[A,B](r)*. Therefore, any positive values for, *p[A,B](r) > p[A]p[B]* indicate a dependence between the points *A* and *B*. Conversely any negative values are indicative of an independent spatial association.

### Modelling microglia fate dynamics

The model for proliferation and fate of microglia distinguishes between proliferating cells (P), which will divide at some point, and non-proliferating cells (N), which have exited cell cycle or are about to do so. When proliferating cells divide, each daughter cell can either go on to proliferate (remaining in state P) or exit cell cycle (attaining state N). Hence, upon division, the following fates are possible:

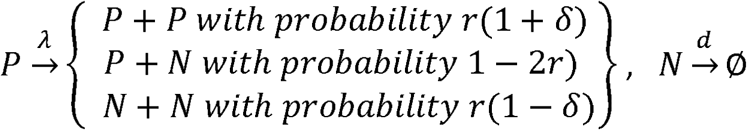

where λ is the division rate of P-cells (here, rates are generally defined as the inverse of the mean time between events). The cell fate probabilities are parametrised by r, the proportion of symmetric divisions, and δ, the bias towards proliferation. Furthermore, we will consider the scenario that P-cells may remain initially quiescent and activate proliferation at a certain time point *t*_0_, which we refer to as model version 2. We also consider the situation where some clones have higher proliferative activity than others, i.e. where λ, r, d or δ variate between clones, (i.e. the proliferative potential is inherited to daughter-P-cells). We implement cell divisions as a Markov process, i.e. cell divisions occur independently of each other at constant stochastic rate. We simulate this model by a Gillespie algorithm (Gillespie, 1977). We note that while the Markov property does not reflect the realistic distributions of individual cell division times, the clonal statistics produced by any cell cycle distribution with a finite variance converge quickly – after a few cell divisions – to the same clonal distribution (this is an instance of a more generalised central limit theorem (Parigini and Greulich, 2020b). Hence a Markov model has the same predictions as a model with more realistic cell cycle time distributions, and we chose the Markov model for its simplicity.

To test the model versions on the NND (nearest neighbour distance) data we used an analytical approach to predict the NND as function of clone size. The NND can be estimated by the average distance, a, between neighbouring cells in a clone, which is related to the cell density of a clone as *a* ∝ *ρ*^−1/3^, with cell density 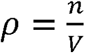, where *n* is the number of cells in a clone and *V* the typical ‘volume’ of the clone. *V* is initially the volume of one cell and grows while the clone spreads at rate 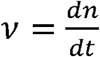 when cells of the clone and other, interspersed cells in the background (belonging to other clones) divide, which occurs at the same rate as the net proliferation rate averaged over all clones, 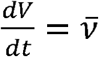. In model version 3, when microglia start proliferating at the same time but with different proliferation rates *v, V* is on average the same for all clones (since the average 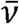 is taken over all clones), and therefore 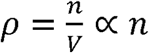, thus *α* ∝ *n*^−1/3^. On the other hand for model version 2, when clones start to proliferate at different times, but the same proliferation rate, 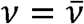, then for each clone both *n* and *V* grow with the same rate, namely 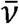, and therefore 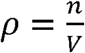 is constant. Hence, also *a* does not vary with *n*. For the latter estimate, we assume that the differences in clone sizes are mainly due to differences in the start time of proliferation rather than stochastic fluctuations (as has been argued in detail in (Klein et al., 2010)).

### Statistical analyses

All data were analysed using the statistical package Graph Pad Prism 9 (GraphPad Software, Inc.). When assumptions of normality were reached, statistical analyses were performed using either a Student’s t-test when comparing between two groups, or a two-way analysis of variance (ANOVA) when comparing across two variables, followed by Tukey’s or Bonferroni’s post-hoc tests. Where assumptions of normality were not reached, either the Kruskal-Wallis or Mann Whitney test were applied. Linear regression and Pearson’s correlation test were performed for correlation analysis. A p value of less than 0.05 was considered statistically significant. Data are expressed as mean ± standard error of the mean (SEM).

