## Supplementary figures and images for "Microglial colonisation of the developing brain is facilitated by clonal expansion of highly proliferative progenitors and follows an allometric scaling"

### S Figure 1

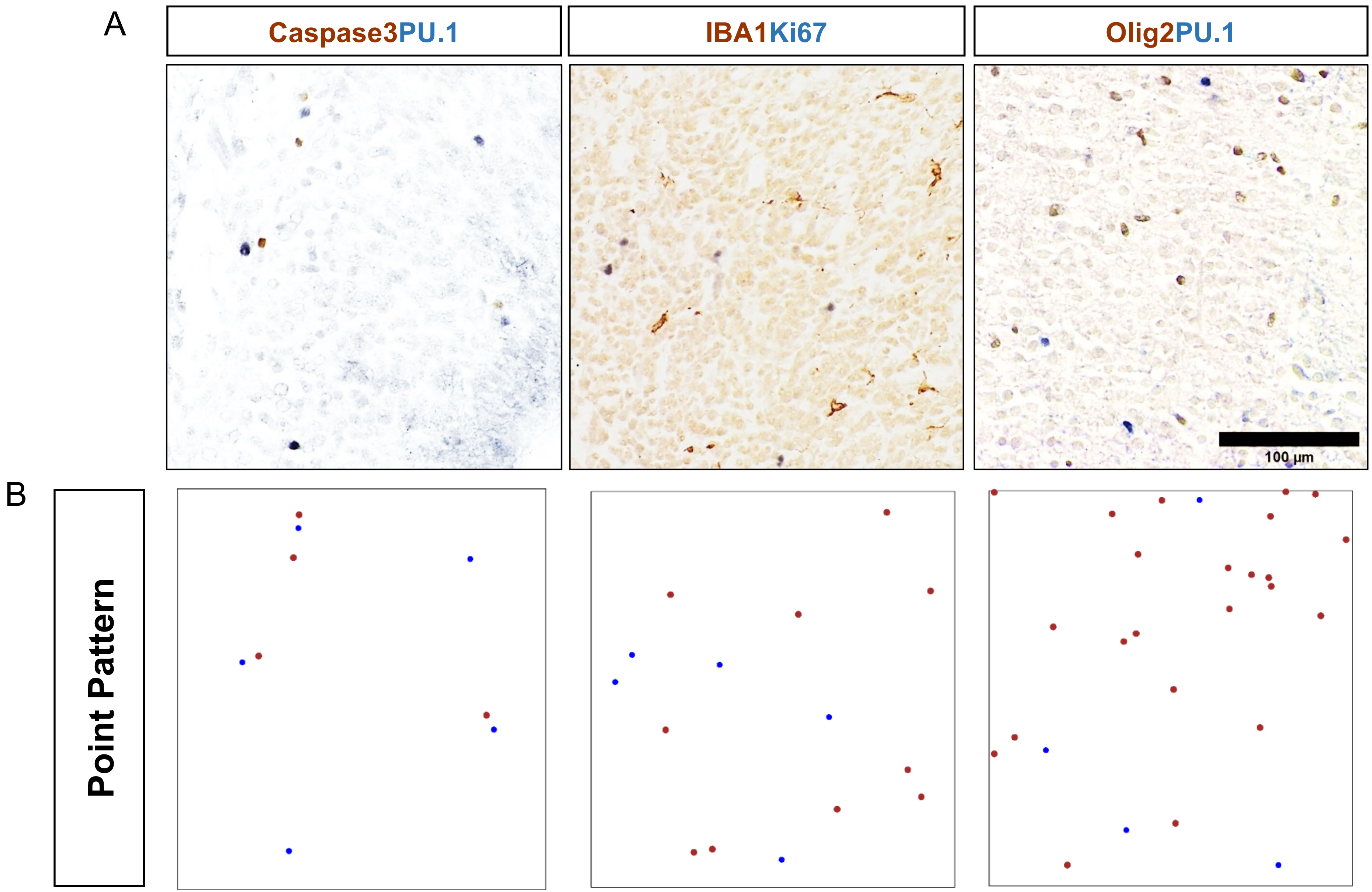

### S Figure 2

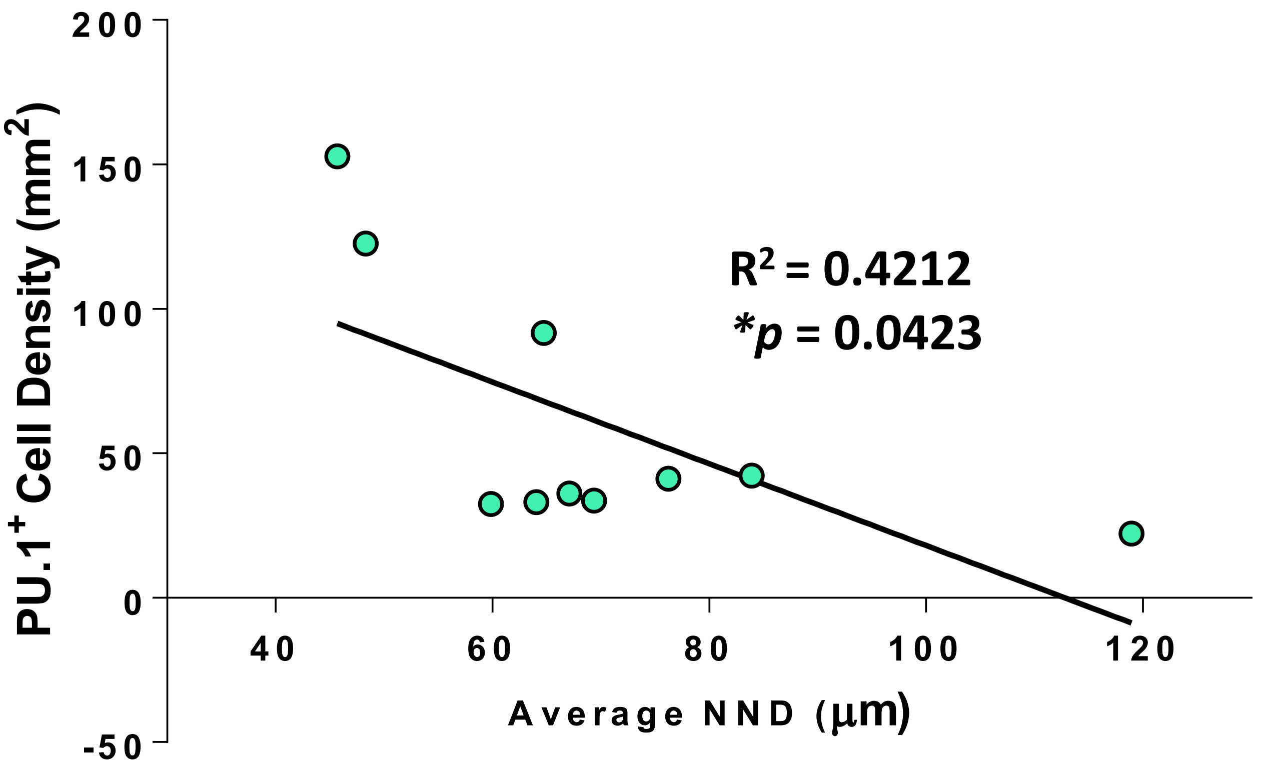

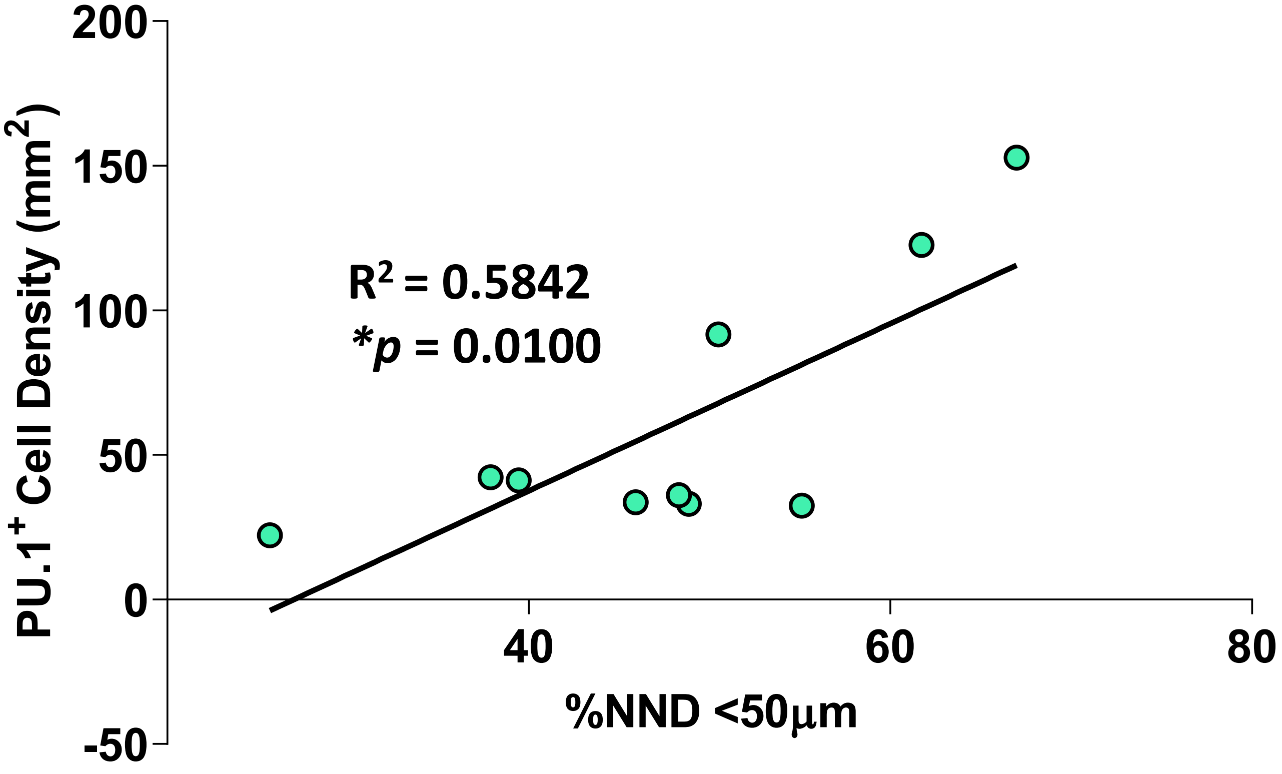

### S Figure 3

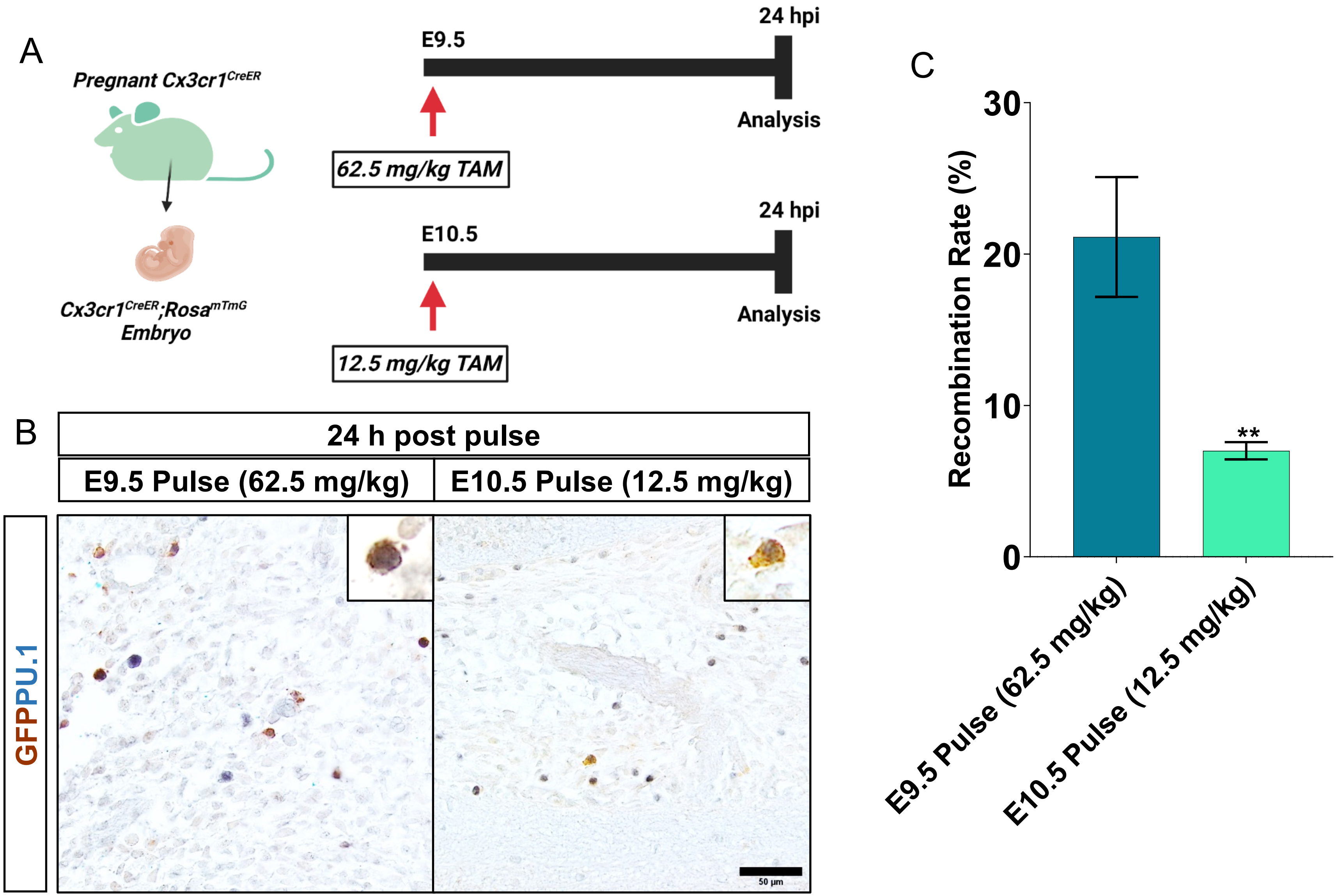

### S Figure 4

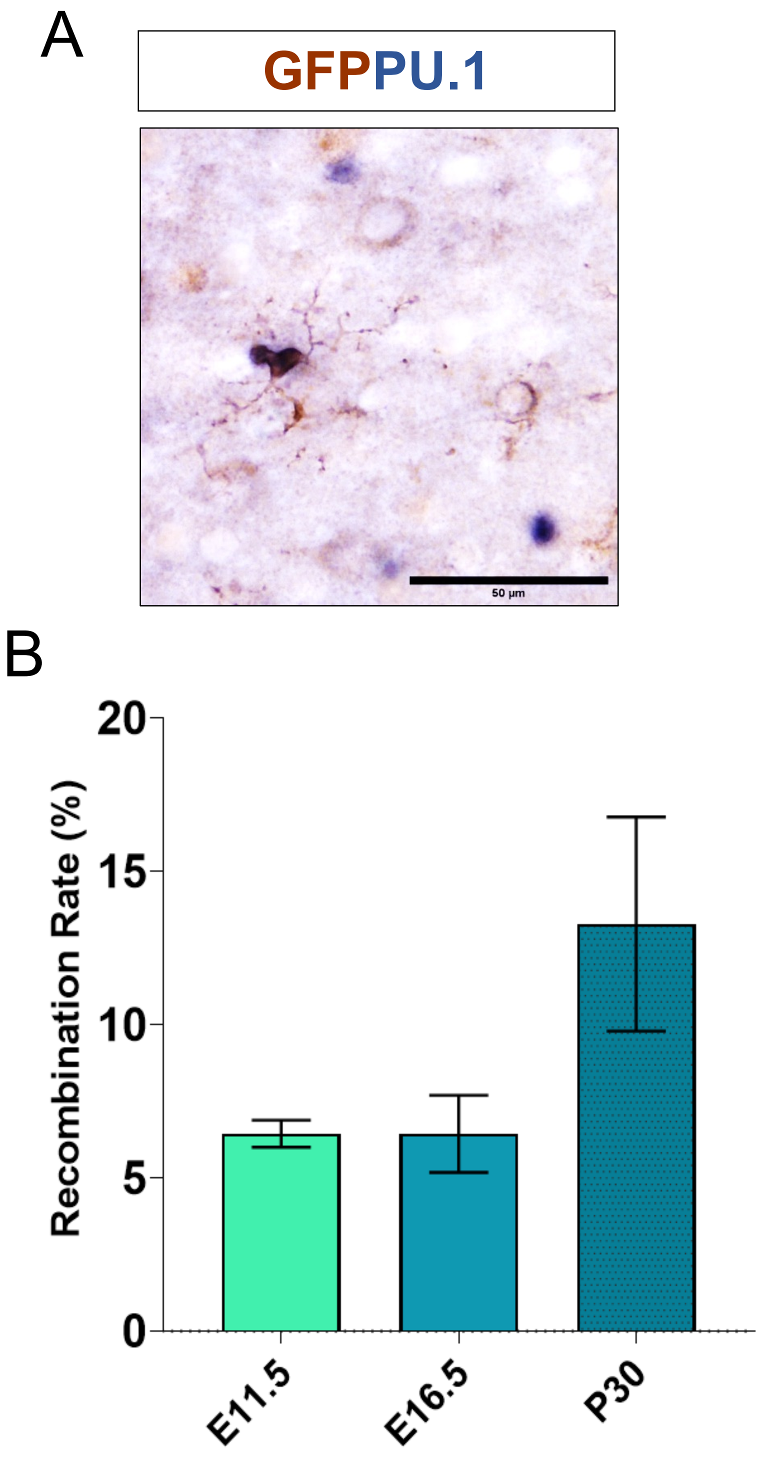

### S Figure 5

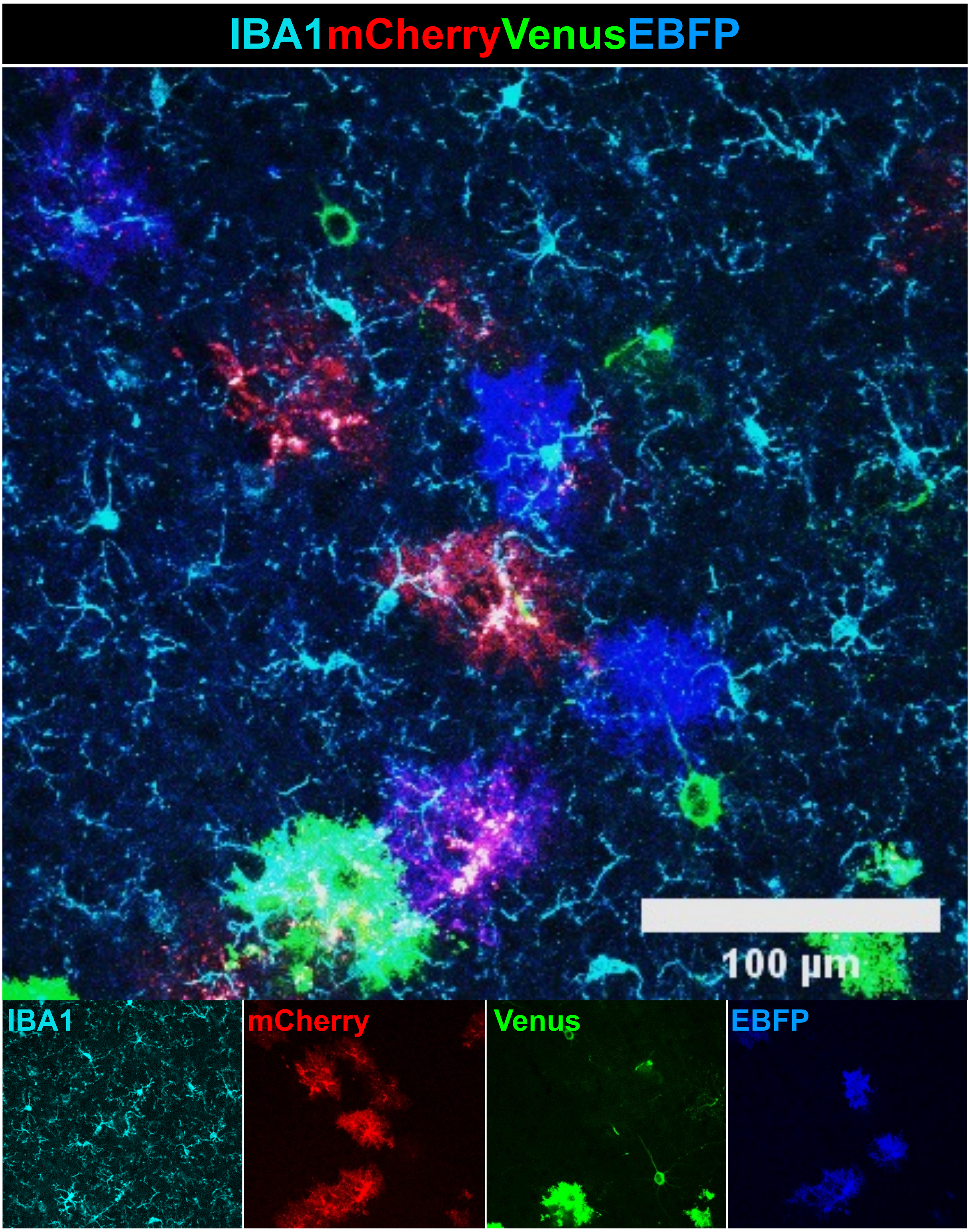

### S Figure 6

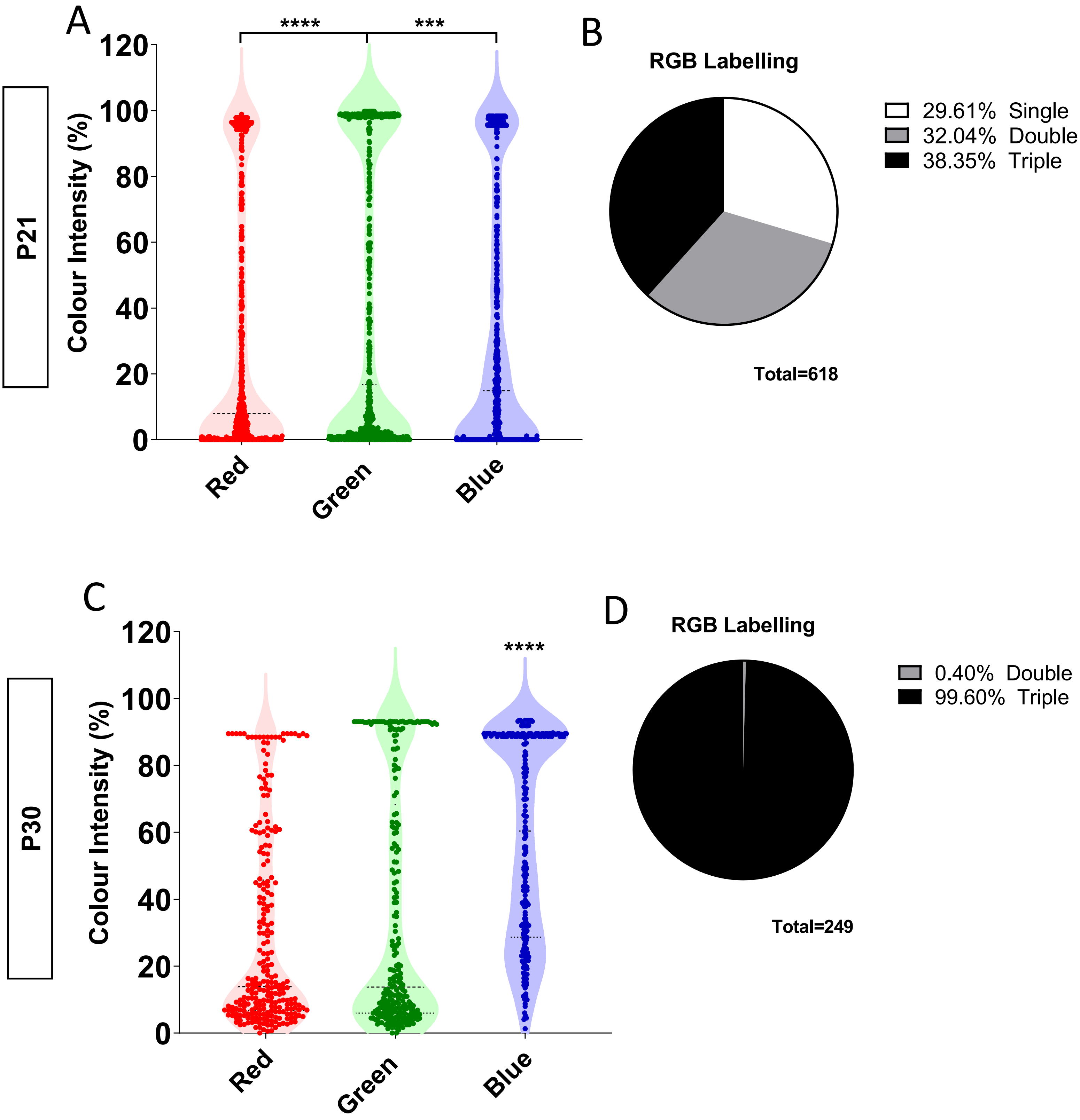

### S Figure 7

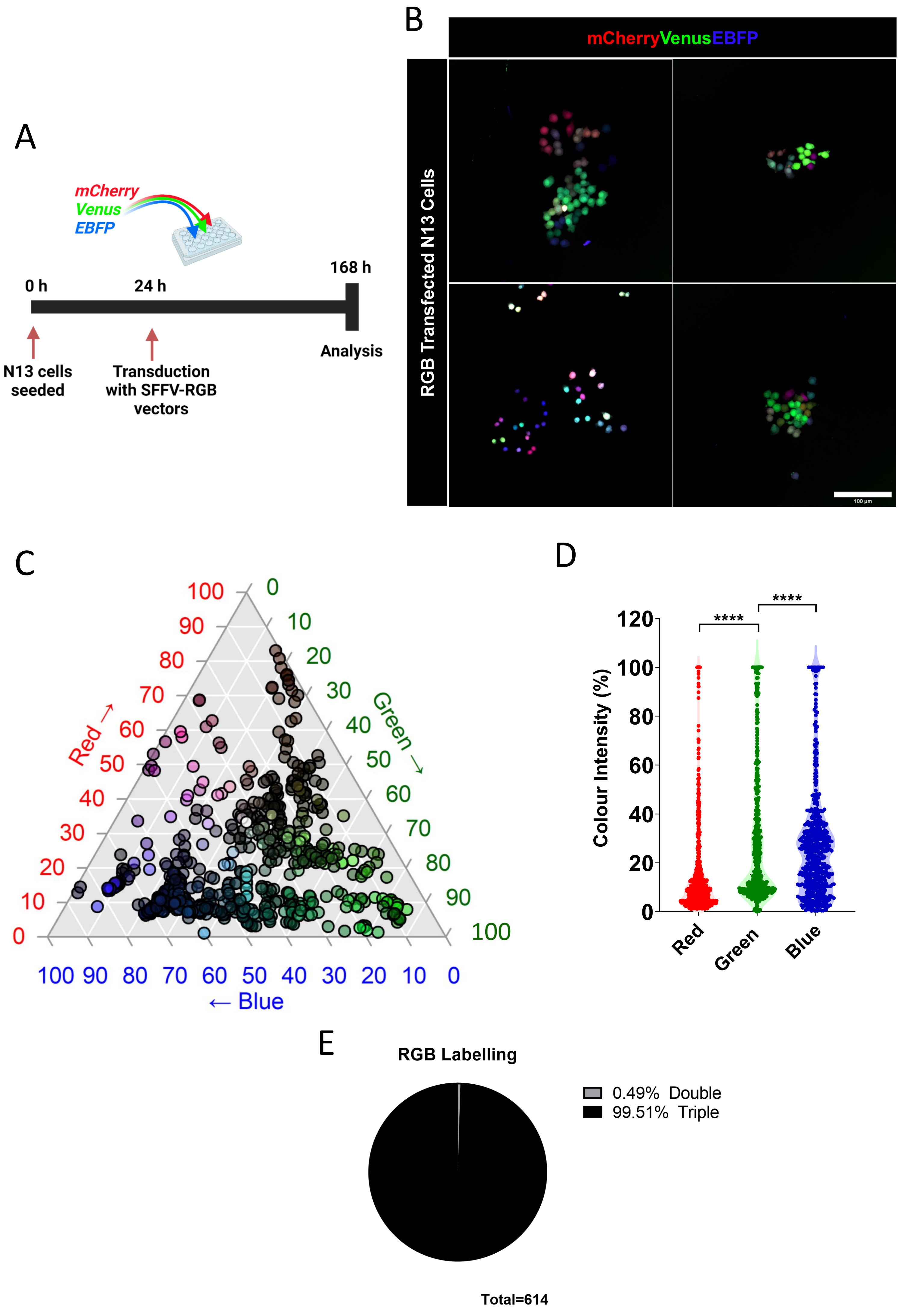

### S Figure 8

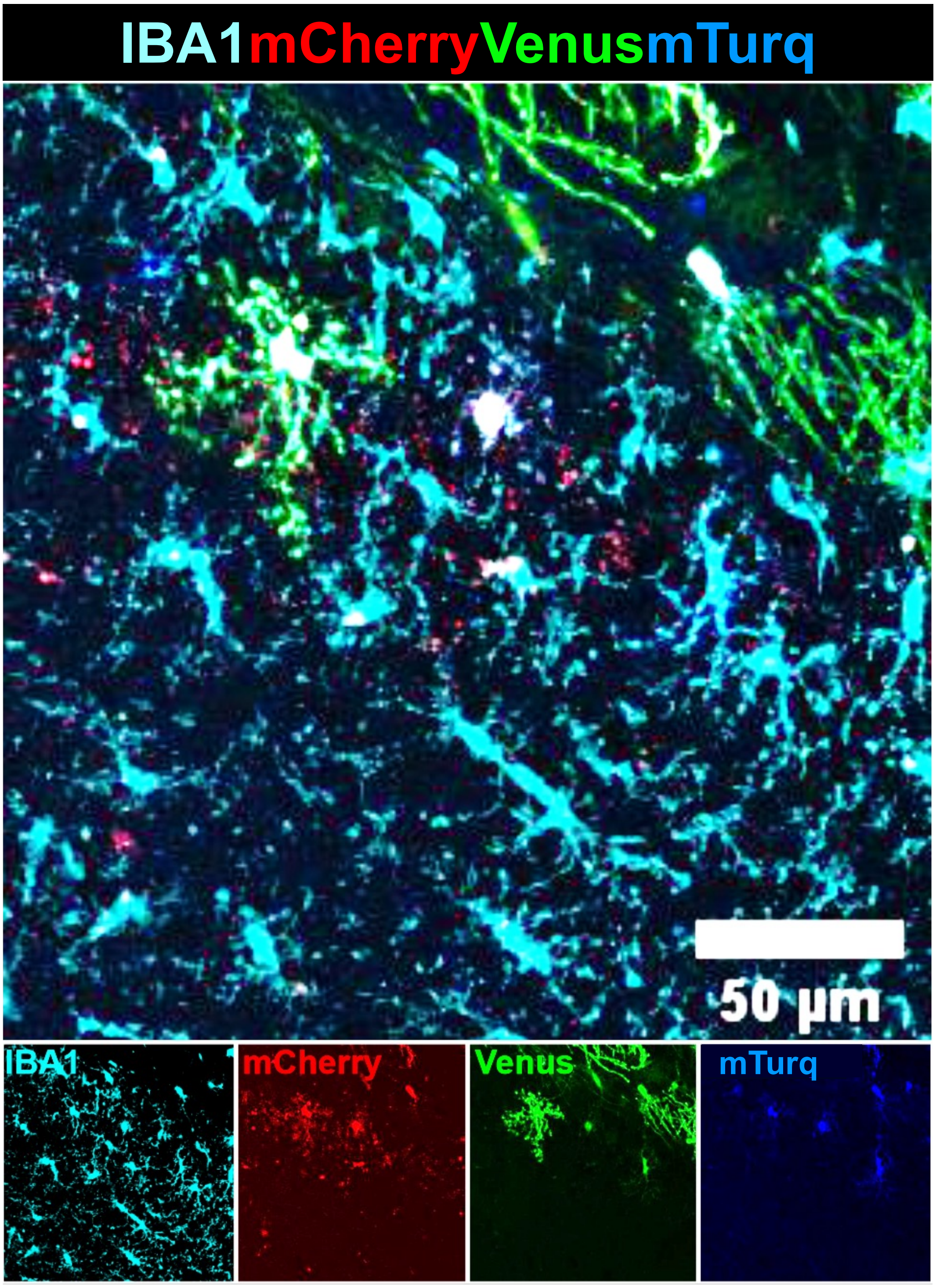
